# Knock-sideways by inducible ER retrieval enables a novel approach for studying *Plasmodium* secreted proteins

**DOI:** 10.1101/2022.10.02.510311

**Authors:** Manuel A Fierro, Tahir Hussain, Liam J Campin, Josh R Beck

## Abstract

Malaria parasites uniquely depend on protein secretion for their obligate intracellular lifestyle but approaches for dissecting *Plasmodium* secreted protein functions are limited. We report knockER, a novel DiCre-mediated knock-sideways approach to sequester secreted proteins in the ER by inducible fusion with a KDEL ER-retrieval sequence. We show conditional ER sequestration of diverse proteins is not generally toxic, enabling loss-of-function studies. We employed knockER in multiple *Plasmodium* species to interrogate the trafficking, topology and function of an assortment of proteins that traverse the secretory pathway to diverse compartments including the apicoplast (ClpB1), rhoptries (RON6), dense granules and parasitophorous vacuole (EXP2, PTEX150, HSP101). Taking advantage of the unique ability to redistribute secreted proteins from their terminal destination to the ER, we reveal vacuolar levels of the PTEX translocon component HSP101 but not PTEX150 are maintained in excess of what is required to sustain effector protein export into the erythrocyte. Intriguingly, vacuole depletion of HSP101 hypersensitized parasites to a destabilization tag that inhibits HSP101-PTEX complex formation but not to translational knockdown of the entire HSP101 pool, illustrating how redistribution of a target protein by knockER can be used to query function in a compartment-specific manner. Collectively, our results establish knockER as a novel tool for dissecting secreted protein function with sub-compartmental resolution that should be widely amenable to genetically tractable eukaryotes.

**Significance:** Protein trafficking and secretion through the endomembrane system is a defining feature of eukaryotes. The secretory pathway is central to the unique biology and pathology of the obligate intracellular malaria parasite, however tools for studying secreted protein function are limited. Knock-sideways is a powerful mutagenesis strategy that conditionally sequesters a protein away from its site of function but is generally not applicable to secreted proteins. We developed a simple approach to conditionally sequester *Plasmodium* secreted proteins in the ER by inducible C-terminal fusion with a KDEL ER-retrieval sequence that can be used for trafficking, topology and loss-of-function studies. The knockER strategy is broadly applicable to functional dissection of proteins that traverse the eukaryotic secretory pathway.

## Introduction

Protein secretion is a fundamental biological process of all living cells. In eukaryotes this is accomplished by a dedicated secretory pathway that begins with import into the Endoplasmic Reticulum (ER) for initial folding and post-translational modification, followed by sorting in the Golgi and post-Golgi compartments. As an obligate intracellular parasite, secreted proteins play a critical role in the biology of the malaria parasite, *Plasmodium*. Through the secretory pathway, proteins are trafficked into both the plasma membrane and several organelles, including a relict plastid called the apicoplast that is critical for parasite metabolism (1, 2). During schizogony, another subset of proteins is packaged into several distinct secretory organelles that are subsequently discharged to facilitate host cell attachment and penetration (3). Invagination of the host membrane by the parasite during invasion generates an intracellular microenvironment called the parasitophorous vacuole (PV) which forms the interface for host-parasite interactions (4). Resident PV proteins secreted into the vacuole perform critical functions at this interface, including nutrient/waste exchange and protein export (5). The latter process enables parasites to deploy a battery of secreted effector proteins across the PV membrane (PVM) and into the host compartment, dramatically remodeling the erythrocyte to meet nutritional demands and avoid host defenses (6, 7). Remarkably, it is estimated that up to 10% of the parasite genome is devoted to encoding exported effectors, further highlighting the central importance of protein secretion to this parasitic lifestyle (8–12).

As with other eukaryotes, entry into the malaria parasite secretory pathway generally begins with co-translational import of proteins containing a signal peptide or transmembrane domain(s) into the ER via the Sec61 complex, followed by sorting from the Golgi to internal compartments, the PV or beyond into the host (13). ER-resident proteins that escape the ER via bulk flow in COPII vesicles are returned to the ER in COPI vesicles by ERD2 surveillance, which binds a C-terminal retrieval sequence (“KDEL” and variants) in the *cis* Golgi and releases it in the ER in a pH-dependent manner (14–16). Similar to other eukaryotes, ER retention of secreted reporters or chimeric proteins can be achieved in *Plasmodium spp.* by a C-terminal KDEL or functional variants (17–22).

In this present work, we developed and characterized a novel DiCre-based knock-sideways tool called knockER that enables conditional fusion of KDEL to the C-terminus of secreted proteins, subjecting the target protein to ERD2 surveillance in the *cis* Golgi to cause its retrieval to the ER. Using this approach, we show successful conditional retrieval of five endogenous *Plasmodium* proteins (ClpB1, RON6, EXP2, PTEX150 and HSP101) that traffic through the ER to diverse compartments including the apicoplast, rhoptries, dense granules, and the PV. Importantly, conditional ER retrieval of both highly expressed reporters and endogenously tagged proteins does not generally impact fitness, indicating knockER-mediated phenotypes are not the indirect result of an integrated stress response, thus making this system suitable for protein loss-of-function studies. Taking advantage of the unique ability to redistribute secreted proteins from their terminal destination to the ER, we reveal a surprising indifference to ER retrieval of HSP101 in *P. falciparum*, in contrast to the other PTEX components, indicating HSP101 is maintained in excess in the PV. Interestingly, redistribution of HSP101 from the PV to the ER hypersensitized parasites to a post-translational destabilization tag that disrupts HSP101 interaction with PTEX but not to translational knockdown, providing support for an HSP101 function in the early secretory pathway in addition to its role in protein translocation across the PVM in the assembled PTEX complex, as recently proposed (23–25). These results establish knockER as a novel tool to study eukaryotic secreted proteins.

## Results

### Conditional ER retrieval of a secreted reporter does not impact parasite fitness

To facilitate conditional ER retrieval of a protein of interest, we designed a strategy based on the dimerizable Cre recombinase (DiCre) system (26, 27) where a gene of interest is C-terminally fused to a 3xFLAG tag and a stop codon flanked by *loxP* sequences imbedded in small introns (28). Activation of DiCre by rapamycin results in excision of the loxP-intervening sequence, removing the 3xFLAG tag and stop codon and bringing into frame a 3xHA tag followed by the ER-retention sequence KDEL and a stop codon (Figure 1A). While the ER plays a crucial role in quality control and is sensitive to added stress, several previous studies have exogenously expressed various KDEL or SDEL fusion proteins in *P. falciparum* and *P. berghei* without apparent impact on parasite fitness (19, 29–36). To directly ascertain whether ER retrieval of a highly expressed secreted protein impacted parasite fitness, we first appended the knockER system to an mNeonGreen (mNG) reporter and fused this to the *exp2* 5’ UTR and signal peptide, enabling expression from the strong *exp2* promoter and entry into the ER followed by default secretion into the PV (37–39). This assembly was placed under control of the anhydrotetracycline (aTc)-responsive TetR-DOZI-aptamers system to prevent expression until aTc is present in the culture (Figure 1A) (40, 41). To combine the Bxb1 integrase system with DiCre, we first introduced a DiCre expression cassette at the benign *pfs47* locus (27) in the *P. falciparum* NF54^attB^ strain (42) (Figure S1) and then integrated the mNG-knockER reporter construct at the *attB* site in the *cg6* locus on chromosome 6 in this NF54^attB-DiCre^ line.

**Figure 1.**
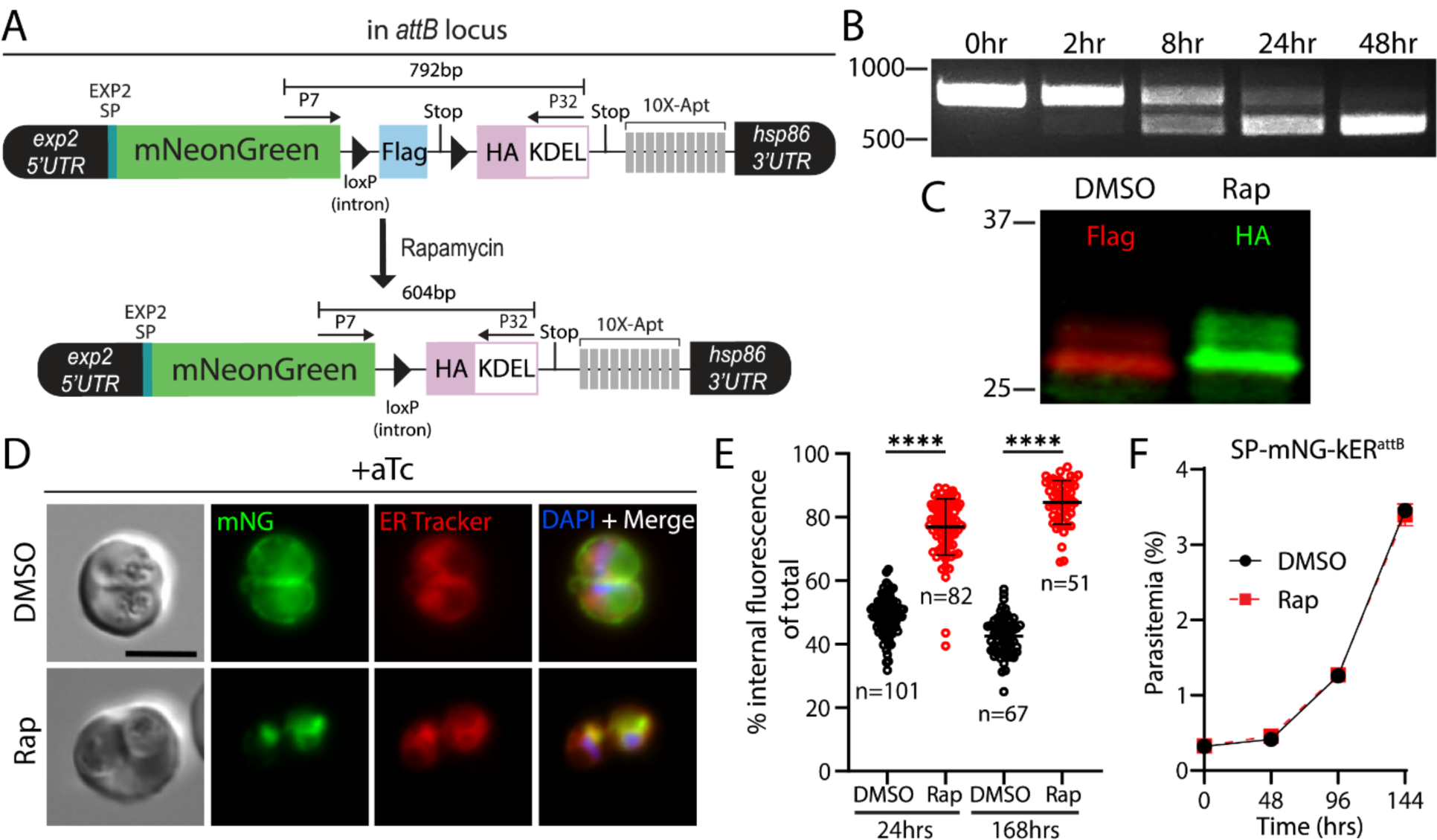
Conditional ER retrieval of a secreted reporter does not impact parasite fitness. A) Schematic of reporter expression cassette installed at the *attB* site of chromosome 6. An mNeonGreen reporter is fused to the signal peptide of EXP2 and expressed under control of the *exp2* promoter. A 3xFLAG tag and stop codon are flanked by loxP-containing introns (triangles) followed by a 3xHA-KDEL, stop codon, and 10X aptamer array (10X-Apt). Aptamer interaction with TetR-DOZI (expressed from a cassette in the plasmid not shown) prevents translation of the reporter, which is relieved in the presence of anhydrotetracycline (aTc) to enable expression. Activation of DiCre by rapamycin treatment excises the FLAG tag, bringing into frame the HA-KDEL tag. B) Time course of excision following rapamycin treatment detected by PCR using primers P7/32. C) Western blot of mNG reporter 24hrs post treatment with DMSO or rapamycin. Molecular weights after signal peptide cleavage are predicted to be 30.8 kDa for mNG-3xFLAG and 30.9 kDa for mNG-3xHA-KDEL. D) Live microscopy of DMSO or rapamycin-treated parasites 24hrs post-treatment. E) Quantification of percent internal mNG fluorescence 24hrs and 168hrs post-treatment with DMSO or rapamycin. Data are pooled from 2 independent experiments and bar indicates mean (****, P<0.0001; unpaired t test). F) Representative growth of asynchronous parasites (n=2 biological replicates) treated with DMSO or rapamycin. Data are presented as means ± standard deviation from one biological replicate (n = 3 technical replicates). All cultures were maintained in media supplemented with 500 nM aTc. Scale bar, 5μm.

Upon treatment with 10nM rapamycin for 3 hours, parasites efficiently excised the knockER cassette (kER) with essentially complete conversion of the population within one parasite developmental cycle (48 hrs) resulting in the expected tag switching from FLAG to HA (Figure 1B,C). Examination of DMSO vehicle-treated parasites by live microscopy showed a peripheral mNG signal, consistent with secretion to the PV (Figure 1D). In contrast, following treatment with rapamycin the mNG signal was dramatically altered to an intracellular, perinuclear pattern that co-localized with ER Tracker, confirming retention in the ER (Figure 1D). Quantification of the distribution of mNG fluorescence between the parasite periphery and interior showed substantial, stable re-localization following KDEL fusion, indicating robust retrieval by ERD2 (Figure 1E, mean % internal mNG at 24hrs: 48.78 ± 5.8% in DMSO vs 76.87 ± 8.9% in Rap, and at 168hrs: 42.54 ± 6.0% in DMSO vs 84.66 ± 6.9% in Rap). Importantly, ER retention of mNG had no impact on parasite growth (Figure 1F) showing that knockER can successfully retrieve a secreted reporter to the ER without impacting parasite fitness.

### knockER reveals Golgi-dependent trafficking and essential apicoplast functionality of ClpB1

Next, we applied knockER to an endogenous gene for conditional loss-of-function mutagenesis. We choose ClpB1, a poorly characterized, nuclear-encoded AAA+ class 1 Clp/HSP100 chaperone that is targeted to the apicoplast (43). As apicoplast ablation can be chemically rescued with isopentenyl pyrophosphate (IPP) (44), we reasoned that IPP supplementation would provide a simple validation that any loss of parasite fitness is a direct effect of sequestering ClpB1 away from its site of action and not an indirect result of ER stress. ClpB1 contains both a signal peptide for ER entry and a transit peptide for apicoplast targeting and localization of ClpB1 to this organelle was previously confirmed by both microscopy and proteomic analyses (35, 43). Using CRISPR/Cas9 editing, we fused mNG to the C-terminus of ClpB1 followed by the kER cassette (Figure 2A). As expected, IFA demonstrated co-localization with the apicoplast marker ACP1, confirming ClpB1 localization to this organelle (Figure 2B). Treatment with rapamycin resulted in efficient excision (Figure 2C) and switching to the 3xHA-KDEL tag produced a shift in migration by Western blot consistent with an increase in molecular weight of ∼14 kDa in the rapamycin treated samples, indicating that the N-terminal transit peptide (residues 24-152, 13.9 kDa) is no longer cleaved in the ClpB1-KDEL fusion (Figure 2D). Since this maturation event takes place in the apicoplast (45), this suggested that ClpB1 was no longer being trafficked to this organelle. Indeed, live microscopy showed a dramatic re-localization of mNG from puncta to a perinuclear distribution that co-localized with ER Tracker, indicative of successful ER retrieval (Figure 2E).

**Figure 2.**
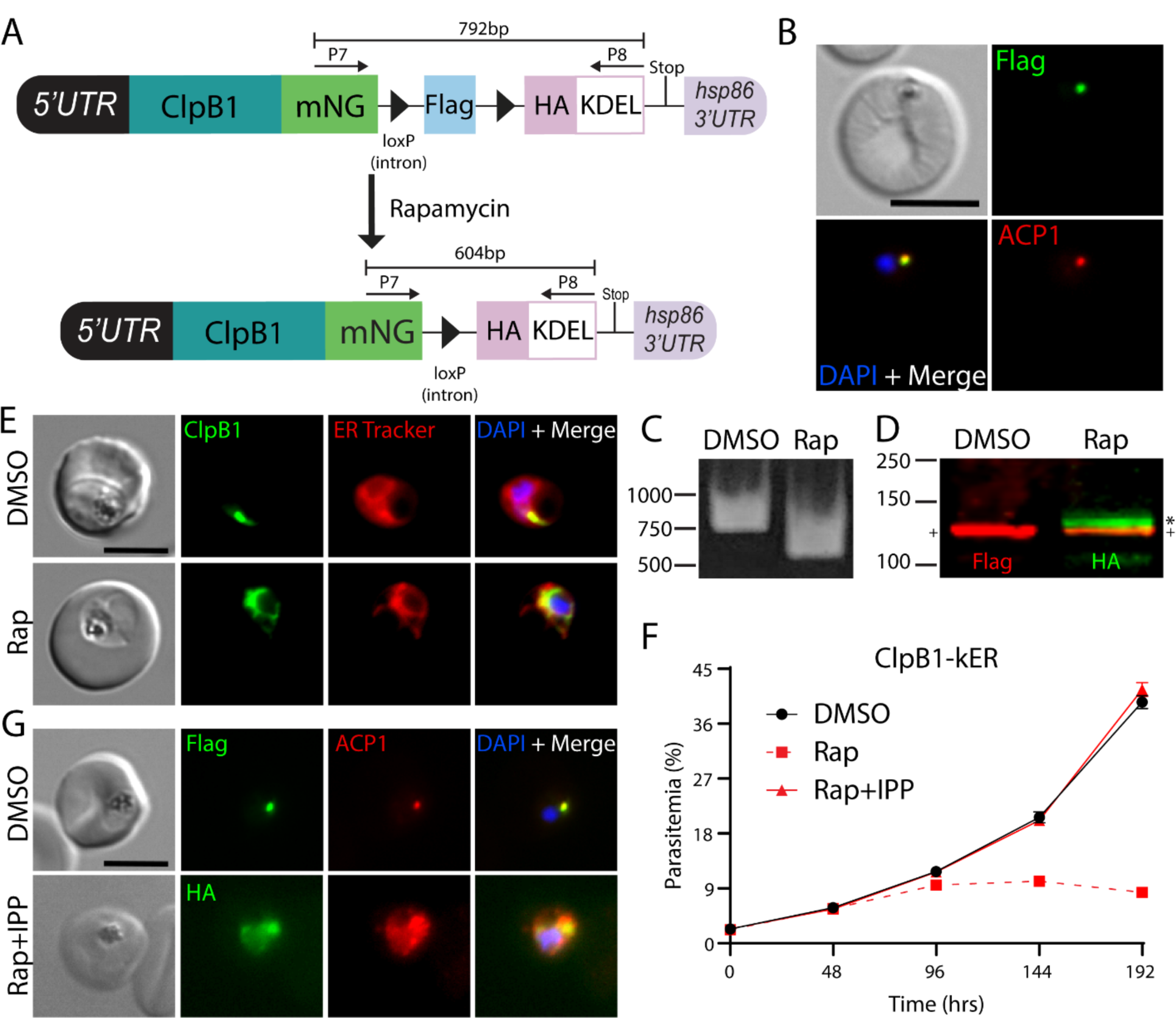
Lethal ER retention of the apicoplast chaperone ClpB1 is rescued by IPP. A) Schematic showing strategy for appending kER to the C-terminus of *clpB1*. B) IFA of PFA fixed parasites stained with mouse anti-Flag and rabbit anti-ACP1 antibodies. C) PCR showing excision in ClpB1-kER parasites 24hrs after rapamycin treatment using primers P7/8. D) Western blot 24hrs post treatment with DMSO or rapamycin. Molecular weights after transit peptide cleavage are predicted to be 135 kDa for ClpB1-mNG-3xFLAG and 135.1 kDa for ClpB1-mNG-3xHA-KDEL. (+) represents the mature form of ClpB1 while the (*) represents ClpB1 with the transit peptide still attached. E) Live microscopy of DMSO or rapamycin-treated parasites 48hrs post-treatment. F) Representative growth of asynchronous parasites (n=3 biological replicates) treated with DMSO or rapamycin, with or without supplementation with 200 µM IPP. Data are presented as means ± standard deviation from one biological replicate (n = 3 technical replicates). G) IFA of PFA fixed parasites stained with mouse anti-Flag or anti-HA and rabbit anti-ACP1 antibodies after 3 growth cycles where rapamycin treated parasites were supplemented with 200 µM IPP. Scale bars, 5μm.

While its functional importance has not been directly tested, ClpB1 is expected to be essential for blood-stage survival based on saturating insertional mutagenesis in *P. falciparum* and pooled knockout screens in *P. berghei* (46, 47). Indeed, ER retention of ClpB1 was lethal and resulted in delayed parasite death (Figure 2F), consistent with an essential role in the apicoplast (48, 49). Moreover, supplementation with IPP fully rescued parasite growth, confirming the observed phenotype was apicoplast-specific and not due to a disruption of ER homeostasis following ClpB1 retention (Figure 2F). Rapamycin-treated parasites supplemented with IPP showed dispersed, vesicular ACP1 signal consistent with plastid disruption, further indicating ClpB1 is required for apicoplast maintenance (Figure 2G). Taken together, these results show for the first time that ClpB1 is an indispensable component of the apicoplast machinery and illustrate that knockER can be used for protein loss-of-function studies in *P. falciparum*.

### knockER induces ER retrieval of rhoptry proteins

Having demonstrated that knockER is suitable to generate loss-of-function phenotypes by protein mis-localization to the ER, we next tested this knock-sideways approach on an expanded set of endogenous soluble and membrane proteins that traffic through the secretory pathway to diverse destinations, including secretory organelles and the PV/PVM. We began by attempting to retrieve RON6, a soluble rhoptry-neck protein that is injected into the PV during invasion (Figure S2A,B). Attempts predating CRISPR technology to disrupt *Pfron6* or truncate its Cysteine-rich C-terminus were unsuccessful (50). More recently, saturating transposon mutagenesis in *P. falciparum* recovered a single insertion within an intron (46) expected to generate a truncation at residue 713 of 950, leaving most of the protein intact (Figure S2A). Curiously, RON6 orthologs in rodent malaria parasites lack the cysteine-rich region which accounts for the C-terminal ∼23% of *P. falciparum* RON6, roughly corresponding to the truncated region in the insertional mutant. As RON6 was not targeted in the PlasmoGEM data set, its suspected essentiality remains uncertain (47). To directly test the importance of RON6, we generated RON6-kER parasites, which underwent efficient excision upon rapamycin treatment (Figure S2C,D). Rapamycin treatment resulted in re-localization of RON6-mNG from the rhoptries to the ER and produced a minor but reproducible growth defect (Figure S2D-F). These results suggest that RON6 may be dispensable in the blood stage, although we cannot exclude the possibility that small amounts of RON6-KDEL may escape ERD2 surveillance at levels sufficient to mediate a critical function. Additionally, the minor fitness defect that results from RON6-KDEL fusion could be the indirect result of modest ER stress induced by RON6 retention. Nonetheless, these findings clearly demonstrate that knockER efficiently retrieves secretory proteins synthesized during the brief period of merozoite formation at the terminal phase of intraerythrocytic development.

### ER retention of EXP2 produces a lethal defect independent to loss of PVM transport function

Having successfully retained several soluble proteins in the ER using knockER, we next tested conditional retrieval of a membrane protein. Solute permeation across the PVM is mediated by a nutrient-permeable channel (51, 52) whose identity was recently tied to EXP2, a PVM protein broadly conserved among vacuole-dwelling apicomplexans (53, 54). EXP2 contains a signal peptide followed by an N-terminal amphipathic helix and oligomerizes to form a heptameric pore in the PVM (25, 55, 56). It is currently unknown how pore formation is specifically constrained to the PVM to avoid perforating other membranes along the secretory pathway or the parasite plasma membrane. While not structurally homologous to the EXP2 pore, bacterial α-helical pore forming toxins are functionally analogous and provide a possible model for EXP2 pore formation. These proteins are typically maintained in a monomeric state that shields a membrane-spanning amphipathic helix, enabling soluble trafficking until contact with components of a target membrane triggers conformational rearrangement, leading to membrane insertion and oligomerization (57). If EXP2 traffics in a similar configuration, then its C-terminus will enter the secretory pathway lumen and be sensitive to ER-retrieval by KDEL fusion. Indeed, we endogenously tagged EXP2 with a version of the kER cassette lacking mNG and found that KDEL fusion disrupted EXP2 trafficking to the PV and resulted in ER retention, confirming its C-terminus resides in the secretory pathway lumen (Figure 3A-D). Notably, ER retrieval of EXP2 appeared to cause more rapid parasite death than previously observed by conditional knockdown or knockout approaches (53, 58, 59) (Figure 3E). To confirm this observation, we engineered parasites to enable DiCre-mediated conditional knockout of EXP2 (EXP2-cKO) to allow for matched induction kinetics to EXP2-kER (Figure S3A-C). Following activation of DiCre in synchronized, ring-stage parasites, the EXP2 knockout progressed to the next cycle with equivalent new ring parasitemia to the DMSO control before death in the second cycle; in contrast, EXP2-kER parasites died immediately in the first cycle and did not form new rings (Figure S3D). To test whether parasite death was solely due to loss of EXP2 transport functions in the PV and not an indirect result of retaining EXP2 in the ER, we generated parasites expressing a second copy of EXP2-mNG-kER from the *attB* locus under the control of its own promoter, leaving the endogenous EXP2 locus unaltered (Figure S3E-G). Retrieval of this second copy of EXP2 to the ER also resulted in parasite death, indicating that EXP2 retention in the ER is toxic independent of compromised transport functions at the PVM (Figure 3F,G). In contrast, ER retrieval of a second copy of EXP2 lacking the amphipathic helix did not cause parasite death (Figure 3H,I and S3E-G). As expected, prior to KDEL fusion the ΔTM version of EXP2 localized to the PV and the digestive vacuole (due to endocytosis of PV proteins along with RBC cytosol). Surprisingly, most cells also showed EXP2ΔTM-mNG export into the RBC cytoplasm (arrow, Figure 3H and S3H). The basis for EXP2ΔTM-mNG export is unclear but may result from interactions with other PTEX components that somehow result in recognition as cargo in the absence of the amphipathic helix. The requirement for the amphipathic helix in EXP2-kER toxicity suggests that increasing EXP2 dwell time in the early secretory pathway may trigger an unfolded protein response or cause aberrant EXP2 pore formation in the ER membrane. Thus, this may provide a new tool for studying EXP2 pore formation but also indicates knockER may not be appropriate for loss-of-function studies of certain membrane proteins.

**Figure 3.**
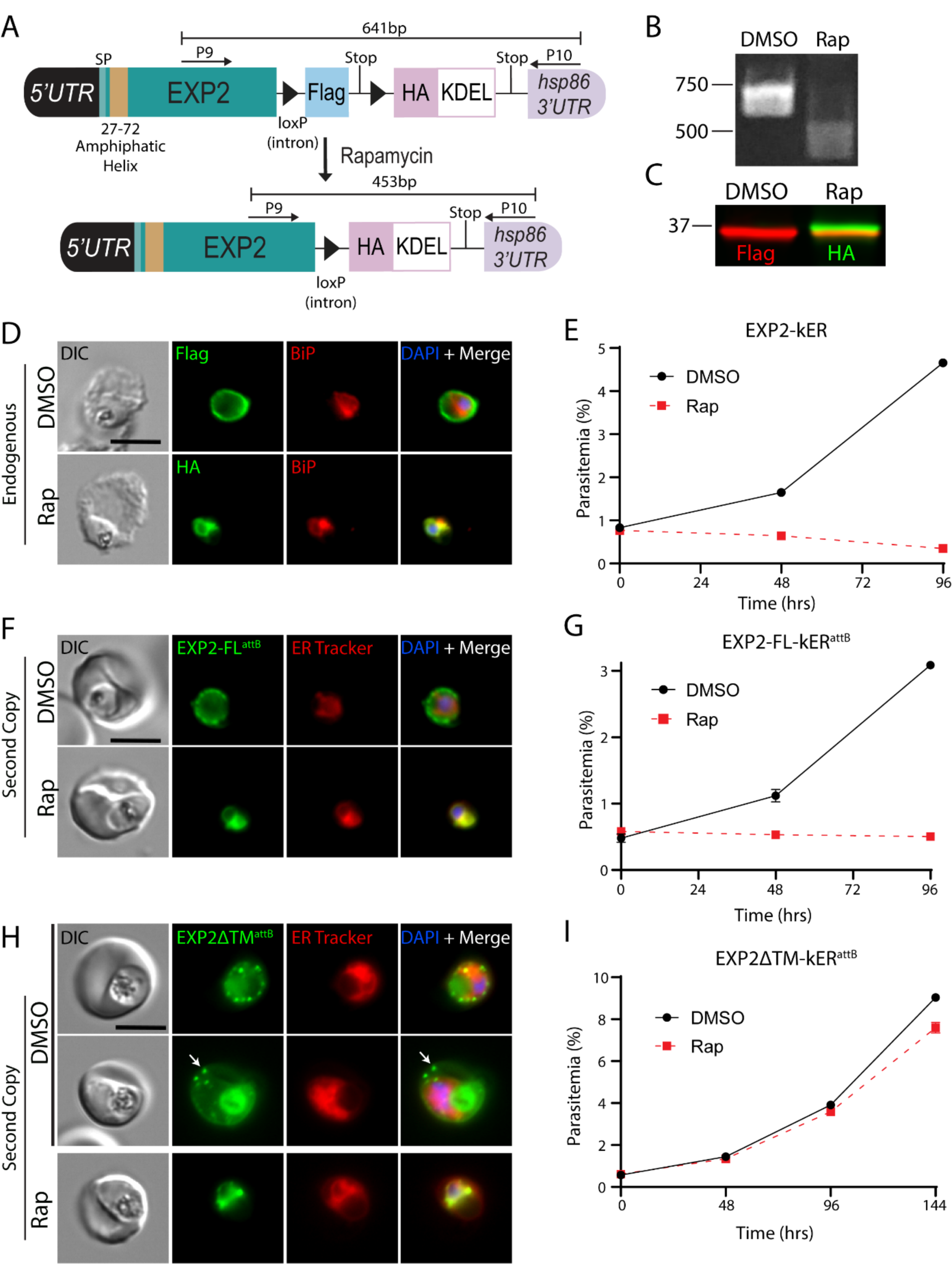
ER retention of EXP2 causes a lethal fitness defect independent of loss of PVM transport functions. A) Schematic showing strategy for appending kER to the endogenous C-terminus of EXP2 without mNG. B) PCR showing excision in EXP2-kER 24hrs after treatment with rapamycin using primers P9/10. C) Western blot of EXP2-kER parasites 24hrs post treatment with DMSO or rapamycin. Molecular weights after signal peptide cleavage are predicted to be 34.6 kDa for EXP2-3xFLAG and 34.7 kDa for EXP2-3xHA-KDEL. D) IFA of EXP2-kER 24hrs post-treatment with DMSO or rapamycin. E) Growth of asynchronous parasites (n=2 biological replicates) treated with DMSO or rapamycin. Data are presented as means ± standard deviation from one biological replicate (n = 3 technical replicates). F) Live microscopy of parasites expressing a full-length second copy of EXP2 with an mNG-kER fusion from the *attB* site under the control of the endogenous *exp2* promoter (EXP2-FL-kER^attB^). Parasites were viewed 24hrs after treatment with DMSO or rapamycin. G) Representative growth of asynchronous EXP2-FL-kER^attB^ parasites (n=2 biological replicates) treated with DMSO or rapamycin. Data are presented as means ± standard deviation from one biological replicate (n = 3 technical replicates). H) Live microscopy of parasites expressing a second copy of EXP2 with mNG-kER fusion and lacking the amphipathic helix from the *attB* site (EXP2ΔTM-kER^attB^). Parasites were viewed 24hrs after treatment with DMSO or rapamycin. Two representative examples of unexcised parasites are presented, showing localization of the ΔTM version of EXP2 to the PV and host cell in some infected RBCs (arrow). Digestive vacuole fluorescence was also observed as is typical for PV proteins. I) Representative growth of asynchronous EXP2ΔTM-kER^attB^ parasites (n=2 biological replicates) treated with DMSO or rapamycin. Data are presented as means ± standard deviation from one biological replicate (n = 3 technical replicates). Second copy EXP2-kER lines were maintained in media supplemented with 500 nM aTc. Scale bars, 5μm.

### ER retention of PTEX150 but not HSP101 results in loss of PTEX function in *P. falciparum*

In malaria parasites, the nutrient-permeable channel has been additionally functionalized by the flange-like adaptor PTEX150 which docks the AAA+ chaperone HSP101 onto EXP2 to form the *<u>P</u>lasmodium* <u>T</u>ranslocon of <u>Ex</u>ported proteins (PTEX), transforming EXP2 into a protein-conducting pore to translocate effector proteins across the PVM and into the erythrocyte (23-25, 53, 60). Interestingly, HSP101 shows a dual localization to both the PV and the ER as opposed to EXP2 and PTEX150, which localize exclusively to the PV and form a sub-complex independent of HSP101 (34, 39, 61, 62). We verified this unique ER localization of HSP101 by protease protection assays in parasite lines expressing fluorescently tagged versions of each PTEX core component, confirming an internal, perinuclear pool of HSP101 but not EXP2 or PTEX150 (Figure S4). This arrangement suggests that in addition to powering translocation in the assembled PTEX complex, HSP101 may also function upstream of the vacuole, possibly initiating cargo selection early in the secretory pathway (61). While several HSP101 conditional knockdown (24, 61) or inactivation (23) strategies have demonstrated an essential role in protein export, it is difficult to resolve distinct compartmental functions with these mutants given that they impact the total protein pool and have no sub-compartmental resolution. In contrast, knockER can potentially resolve functions across distinct compartments by redistributing a target protein pool from its terminal destination to the ER.

In an attempt to separate HSP101 ER function from its role in the PV, we generated a kER fusion to the endogenous copy of HSP101, expecting that redistribution of HSP101 to the ER would be lethal but might provide insight into any ER localized function which could be preserved compared to the phenotype of HSP101 knockdown mutants. In parallel, we also generated a PTEX150-kER line, reasoning that since HSP101 and PTEX150 have similar expression timing and are exclusively involved in protein export but not nutrient uptake (53), PTEX150 would provide a control for knockER inactivation of PTEX activity independent of ER function. As expected, the PTEX150-KDEL fusion was efficiently retrieved to the ER, producing a lethal growth defect and accompanying block in protein export (Figure 4A-E), similar to *glmS* ribozyme-mediated knockdown of PTEX150 (24). Importantly, ER retention of a second copy of PTEX150-kER in parasites where the endogenous PTEX150 locus was unmodified did not impact growth, demonstrating that the phenotype observed upon ER retention of the endogenously tagged protein is a direct result of functional disruption of PTEX150 (Figure S5A-D).

**Figure 4.**
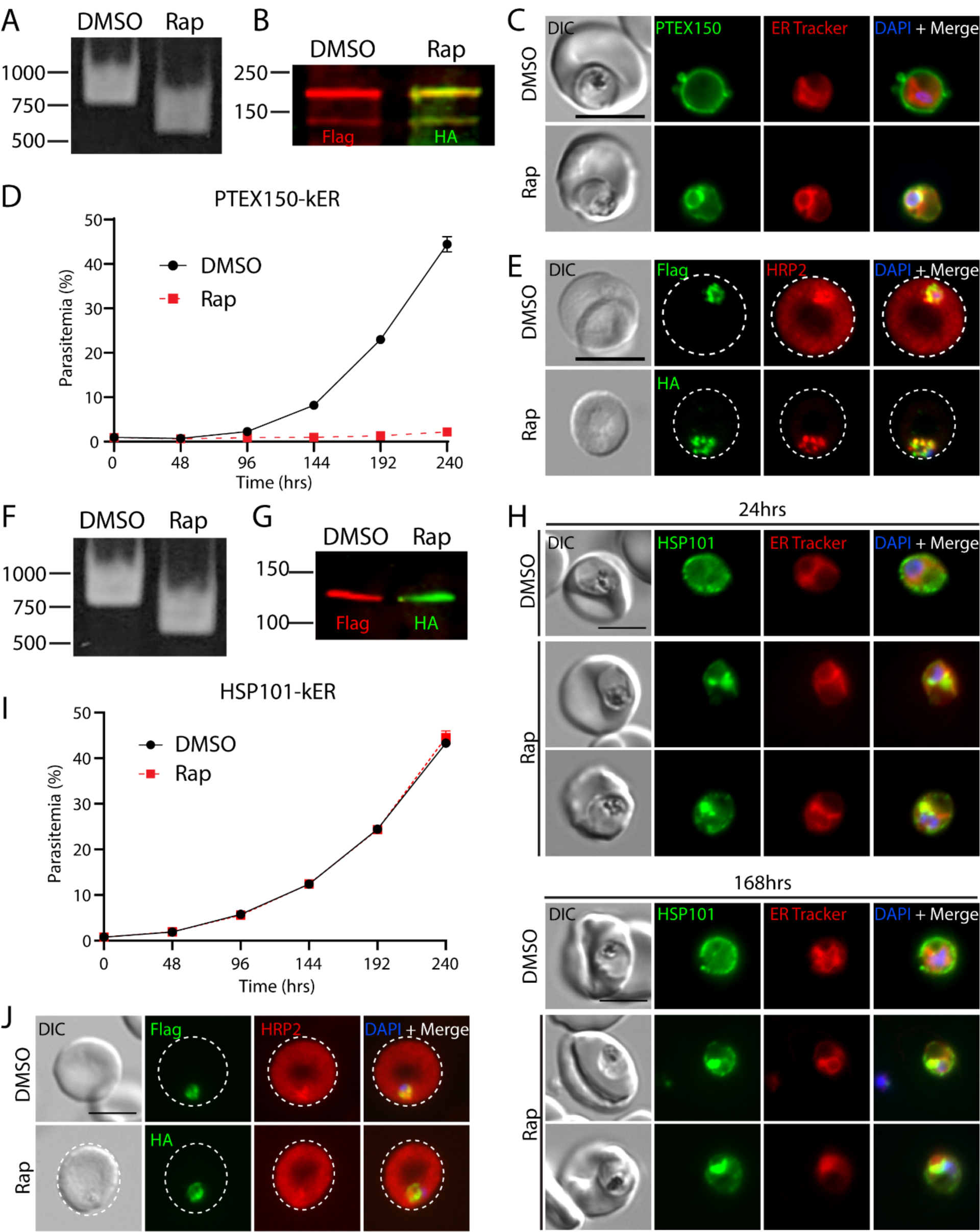
ER retention of PTEX150 but not HSP101 causes a lethal block in protein export in *P. falciparum*. A) PCR showing excision in PTEX150-kER parasites 24hrs after rapamycin treatment using primers P7/8. B) Western blot of PTEX150-kER 24hrs post-treatment with DMSO or rapamycin. Molecular weights after signal peptide cleavage are predicted to be 141 kDa for PTEX150-mNG-3xFLAG and 141.1 kDa for PTEX150-mNG-3xHA-KDEL. Note that PTEX150 is observed to migrate at higher molecular weight than predicted (60). C) Live microscopy of PTEX150-kER 24hrs post-treatment with DMSO or rapamycin. D) Representative growth of asynchronous PTEX150-kER parasites (n=3 biological replicates) treated with DMSO or rapamycin. Data are presented as means ± standard deviation from one biological replicate (n = 3 technical replicates). E) IFA of synchronized, ring-stage PTEX150-kER parasites 48hrs post-treatment with DMSO or rapamycin and probed with anti-HRP2 and anti-FLAG or anti-HA antibodies. F) PCR showing excision in HSP101-kER parasites 24hrs after rapamycin treatment using primers P7/8. G) Western blot of HSP101-kER parasites 14 days post-treatment with DMSO or rapamycin. Molecular weights after signal peptide cleavage are predicted to be 130.5 kDa for HSP101-mNG-3xFLAG and 130.6 kDa for HSP101-mNG-3xHA-KDEL. H) Live microscopy of HSP101-kER 24hrs and 168hrs post-treatment with DMSO or rapamycin. I) Representative growth of asynchronous HSP101-kER parasites (n=3 biological replicates) treated with DMSO or rapamycin. Data are presented as means ± standard deviation from one biological replicate (n = 3 technical replicates). J) IFA of synchronized, ring-stage HSP101-kER parasites 48hrs post-treatment with DMSO or rapamycin and probed with anti-HRP2 and anti-FLAG or anti-HA antibodies. Scale bar, 5μm.

Unexpectedly, although rapamycin-induced HSP101-KDEL fusion increased ER localization similar to PTEX150 (Figure 4F-H), parasite growth was completely unaffected and protein export was not detectibly impacted (Figure 4I,J), sharply contrasting with the lethal export defect observed with HSP101 depletion or inactivation mutants (23, 24, 61). Indeed, HSP101-kER parasites could be cultured indefinitely following excision and displayed sustained ER retention of HSP101 (Figure 4H and S5E,F, mean Rap internal HSP101: 58.82±9.58% at 24 hours vs 54.82±9.84% at 7 days). As an orthogonal approach to monitor protein redistribution by KDEL fusion, we released infected RBCs with saponin to permeabilize the PVM and quantified fluorescence via flow cytometry with or without proteinase K treatment. Again, we observed an increase in protected fluorescence following rapamycin treatment, consistent with redistribution of HSP101 to the ER (Figure S5G). Importantly, PCR and Western blot indicated that rapamycin-treated parasites had undergone complete excision with the pre-excised FLAG-tagged version of HSP101 no longer detectible (Figure 4F,G). Furthermore, both the mNG-FLAG and mNG-HA-KDEL tagged versions of HSP101 migrated as a single band at the expected size for the full-length fusion proteins with no apparent breakdown products, ensuring that the mNG signal distribution observed by live microscopy reflected an intact HSP101 fusion (Figure 4G, S7B and S8B).

While a large fraction of PTEX150 and HSP101 was clearly ER-retained following knockER induction, some peripheral signal was still apparent after rapamycin treatment (Figure 4C,H). To determine whether the observed phenotypic differences between HSP101 and PTEX150 were due to a difference in sensitivity to ERD2 surveillance, we quantified the internal mNG fluorescence signal in HSP101- and PTEX150-kER parasites following KDEL fusion. In synchronized parasites, a similar increase in internal signal was observed in both lines at 24 hours post Rap treatment (Figure S5E, mean Rap internal HSP101 58.82±9.58% vs 56.32±18.56% for PTEX150). Since a comparison could not be made with PTEX150-kER parasites at 7 days due to parasite death by the second cycle following excision, we compared HSP101-kER parasites with second copy PTEX150-kER^attB^ control parasites 7 days after excision and again observed a comparable internal signal redistribution (Figure S5F, mean Rap internal HSP101 54.82±9.84% vs 61.75±16.27% for PTEX150 at 7 days). As major differences in ER retrieval cannot account for parasite insensitivity to HSP101-kER, these results indicate vacuolar HSP101 levels are unexpectedly maintained in excess from what is required for PTEX function at the PVM.

### ER retention of *P. berghei* HSP101 results in a lethal export defect

To determine if ER retention of HSP101 was similarly tolerated in other *Plasmodium* species, we tagged HSP101 and PTEX150 with kER in a *P. berghei* rodent malaria DiCre parasite. We also included a downstream cassette in this tagging plasmid for expression of a secreted mRuby under control of the constitutive *Pbhsp70* promoter to label the PV (Figure S6A). Similar to *P. falciparum* knockER mutants, rapamycin treatment produced efficient excision and ER retrieval of PbPTEX150 and PbHSP101 (Figure 5A-C,F). Curiously, the localization pattern of KDEL fusion proteins in *P. berghei* parasites was clearly internal to PV-mRuby but more diffuse than in *P. falciparum*, suggesting distinctive ER morphology between these two species. Interestingly, PbHSP101-kER and PbPTEX150-kER parasites displayed greater internal retention than the corresponding *P. falciparum* knockER mutants (HSP101: mean % internal mNG 77.42 ± 7.72% in *P. berghei* vs 58.82±9.58% in *P. falciparum*; and PTEX150: mean % internal mNG 69.55 ± 10.86% in *P. berghei* vs 56.32±18.56% in *P. falciparum*), suggesting that KDEL-mediated ER retrieval is more stringent in rodent parasites (Figure 5D,G). To evaluate protein export, we generated versions of the PbPTEX150-kER and PbHSP101-kER mutants where the PV-mRuby reporter was replaced with a cassette for expression of the exported protein IBIS1 fused to mRuby (Figure S6A) (63). In contrast to *P. falciparum*, KDEL fusion to both PbPTEX150 and PbHSP101 produced a stark block in IBIS1 export (Figure 5I,J). To determine if KDEL fusion to PbHSP101 impacted parasite fitness, parasites were induced *ex vivo* followed by overnight culture to allow for tag switching before equal numbers of DMSO or rapamycin treated parasites were IV injected into naïve mice. Strikingly, while DMSO controls became patent by day two post-injection, neither KDEL-fused PbPTEX150 or PbHSP101 parasites appeared over the course of 10 days before the experiment was terminated (Figure 5E,H and S6B).

**Figure 5:**
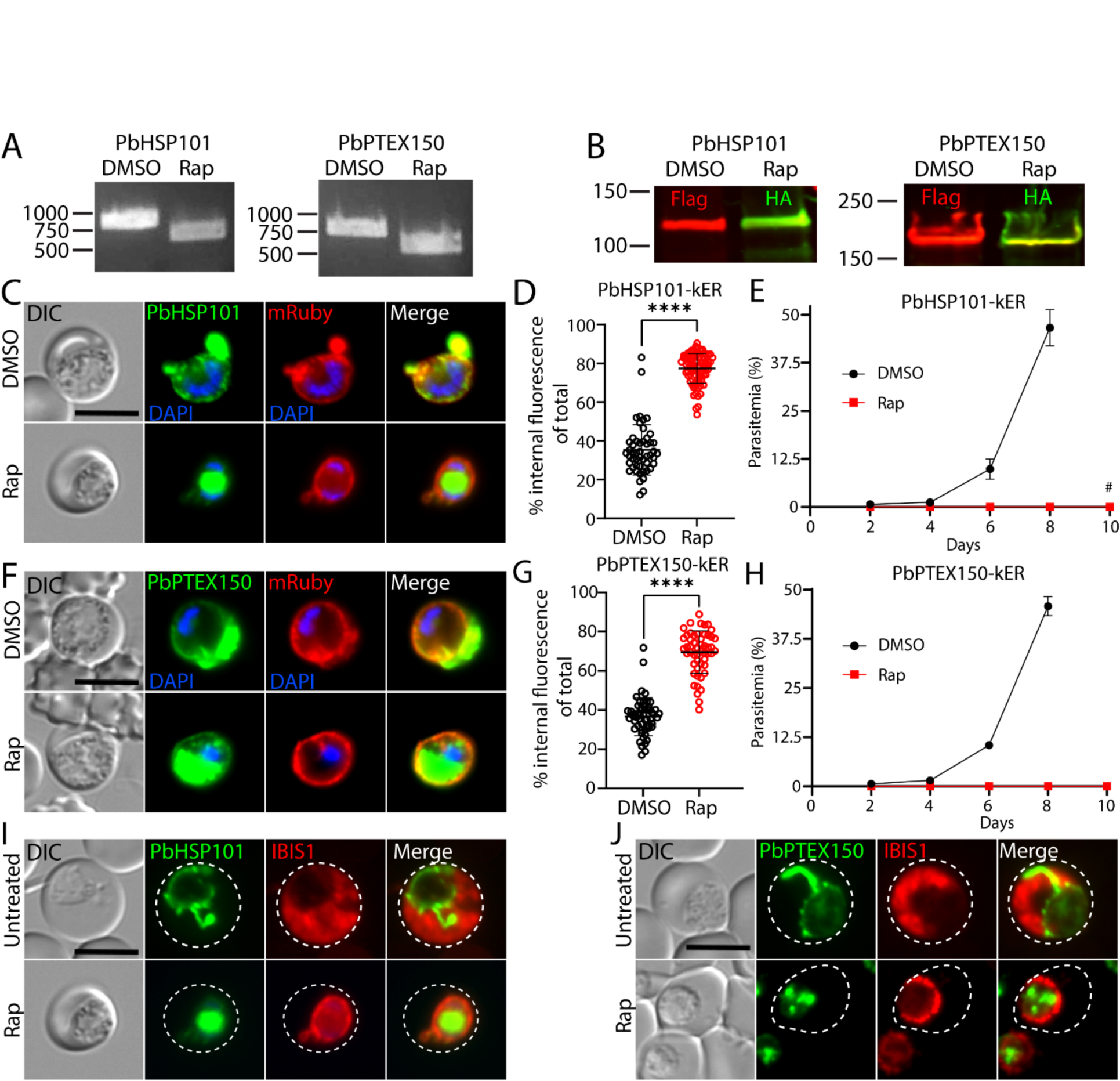
ER retention of HSP101 causes a lethal block in protein export in *P. berghei*. A) PCR showing excision in PbPTEX150-kER and PbHSP101-kER *P. berghei* parasites treated with rapamycin and cultured 18hrs *ex vivo* using primers P7/P60. B) Western blot of PbPTEX150-kER and PbHSP101-kER parasites 24hrs post-treatment with DMSO or rapamycin. Molecular weights after signal peptide cleavage are predicted to be 130.9 kDa for PbHSP101-mNG-3xFLAG, 131 kDa for PbHSP101-mNG-3xHA-KDEL, 130.5 kDa for PbPTEX150-mNG-3xFLAG and 130.6 kDa for PbPTEX150-mNG-3xHA-KDEL. Note that PbPTEX150 is observed to migrate at higher molecular weight than predicted (101). C,F) Live microscopy of PbHSP101-kER or PbPTEX150-kER parasites 18hrs post-treatment with DMSO or rapamycin. These transgenic lines also contain a downstream cassette for expression of mRuby bearing a signal peptide for secretion into the PV (see Figure S6A). D,G) Quantification of percent internal mNG fluorescence 24hrs post-treatment with DMSO or rapamycin. Data are pooled from 2 independent experiments and bar indicates mean (****, P<0.0001; unpaired t test). E,H) Parasitemia of PbHSP101-kER or PbPTEX150-kER infected mice inoculated by tail vein-injection of parasites treated *ex vivo* with DMSO or rapamycin. Data are presented as means ± standard deviation of three independent experiments (one mouse infected per treatment in each experiment). No parasites were observed in mice injected with rapamycin treated parasites except that one mouse injected with rapamycin-treated PbHSP101-kER parasites became patent on day 10 (#). However, PCR showed these parasites had not undergone excision (see Figure S6B). I,J) Live microscopy of PbHSP101-kER or PbPTEX150-kER parasites containing a downstream cassette for expression of an IBIS1-mRuby fusion that is exported into the RBC (See Figure S6A). Parasites were imaged before or 24hrs after treatment with rapamycin. Dashed lines indicate RBC boundaries traced from DIC images. Images are representative of n=2 biological replicates. Scale bars, 5μm.

The block in IBIS1 export suggests that PbHSP101-KDEL is sufficiently retained to produce a lethal export defect, in contrast with PfHSP101-KDEL; however, phenotypes can be masked by the rich resources available in culture (64, 65) and many exported effectors critical for survival in the vertebrate host are dispensable *in vitro* (12) where parasites do not have to contend with host defenses (66, 67). To distinguish whether the fitness defect incurred by ER retention of PbHSP101 resulted simply from parasite death or from the unique pressures imposed by the host, we evaluated parasite development of synchronized ring-stage cultures *ex vivo*. Importantly, both PbHSP101-kER and PbPTEX150-kER parasites also failed to develop into terminal schizonts *in vitro* following rapamycin treatment, indicating the *in vivo* fitness defect is not merely the result of host defenses (Figure S6C-E). Taken together, our data show that ER-retention of HSP101-KDEL is more efficient in *P. berghei,* resulting in a lethal defect in protein export.

### ER retention of HSP101 hypersensitizes parasites to a destabilization tag that inhibits HSP101-PTEX complex formation

The lack of an observable phenotype after ER-retention of HSP101 in *P. falciparum* suggests that the level of retention does not reach a critical threshold required to impact HSP101 function at the PV. To determine the amount of HSP101 needed to support *P. falciparum* growth *in vitro*, we combined knockER with the TetR-DOZI-aptamers system to generate a HSP101-kER line that also allows for titration of HSP101 expression with aTc (Figure 6A). Like the original HSP101-kER line, rapamycin-treated HSP101-kER^TetR-DOZI^ parasites exhibited ER retention of HSP101 without impacting parasite growth or export of representative PEXEL and PNEP proteins (Figure 6B-D). However, HSP101 depletion following aTc washout resulted in a robust block in protein export and parasite death (Figure 6D and S7A,B), consistent with previous HSP101 knockdown or inactivation studies (23, 24, 61). To determine the level of total HSP101 reduction sufficient to impact growth, we grew synchronized parasites at a range of aTc concentrations following DMSO or rapamycin treatment and measured mNG fluorescence via flow cytometry in parallel with parasite growth (Figure 6E,F). Interestingly, rapamycin-treated parasites grown at higher aTc concentrations displayed a small but reproducible increase in total HSP101, consistent with a reduction in HSP101 turnover by endocytosis from the PV to the digestive vacuole following redistribution to the ER (Figure 6E). More importantly, we determined that a ∼70% reduction in total HSP101 levels was necessary to impact parasite growth, showing that a relatively small amount of HSP101 is needed for survival (Figure 6E,F, 5nM aTc), consistent with redistribution of HSP101 by knockER not meeting the threshold required to impact HSP101 PV function and parasite fitness in *P. falciparum*.

**Figure 6.**
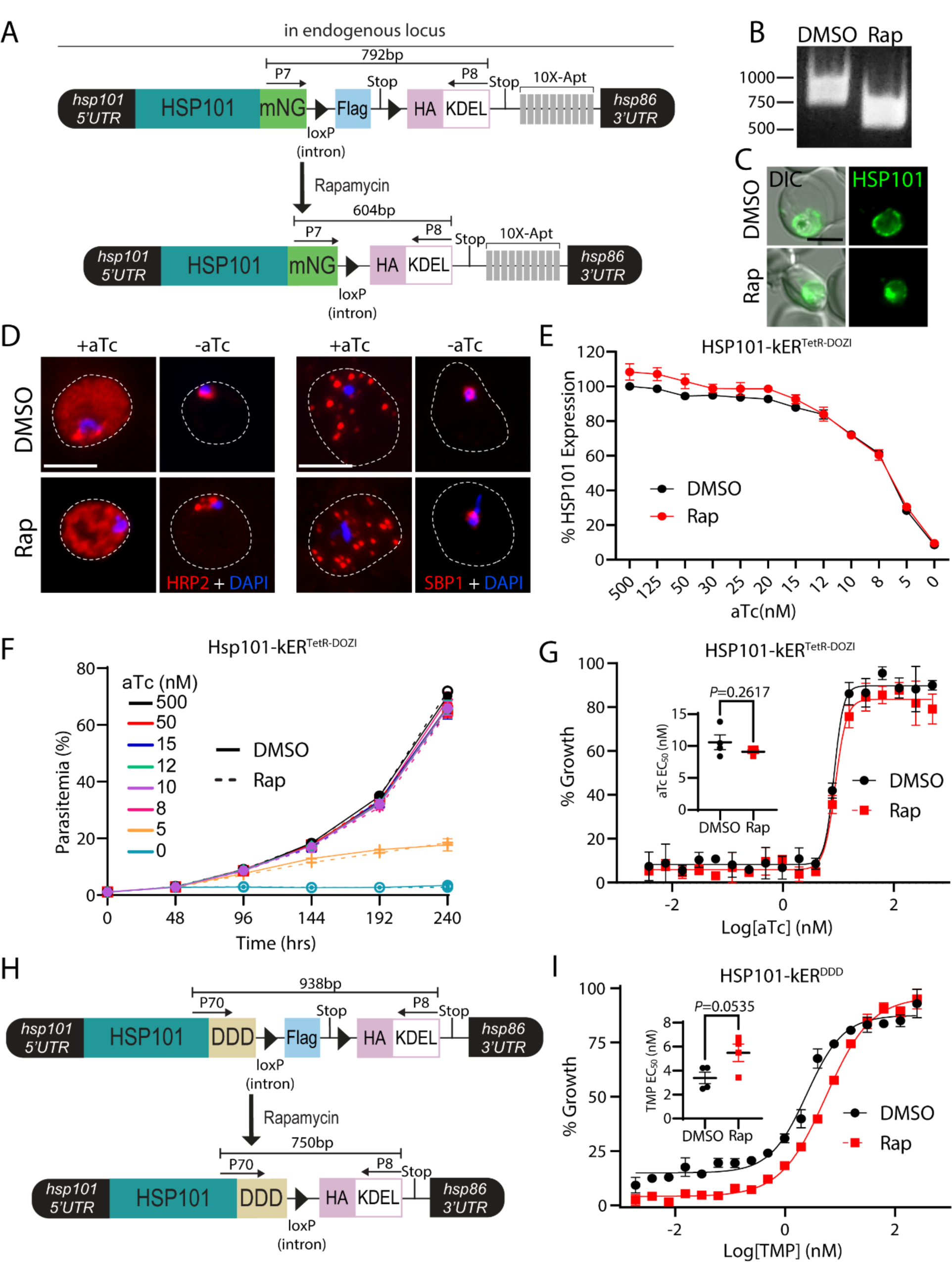
Depletion of HSP101 PV levels hypersensitizes parasites to post-translational destabilization of the HSP101-PTEX complex but not HSP101 translational knockdown. **A)** Schematic showing strategy for appending kER and the TetR-DOZI aptamers system to the endogenous C-terminus of HSP101. B) PCR showing excision in HSP101-kER^TetR-DOZI^ parasites 24hrs after rapamycin treatment using primers P7/8. C) Live microscopy of HSP101-kER^TetR-DOZI^ parasites 168hrs post-treatment with DMSO or rapamycin. D) IFA detecting the soluble PEXEL protein HRP2 or the transmembrane PNEP SBP1 in synchronized, ring-stage HSP101-kER^TetR-DOZI^ parasites grown for 48hrs +/-aTc and treated with DMSO or rapamycin 1 cycle prior. Dashed lines indicate RBC boundaries traced from DIC images. E) Quantification of HSP101 levels by flow cytometry measurement of mNeonGreen fluorescence in HSP101-kER^TetR-DOZI^ parasites grown for 48 hrs at indicated aTc concentrations and treated with DMSO or rapamycin 2 cycles prior. MFI data were normalized to DMSO-treated parasites grown in 500 nM aTc (100%) and presented as mean ± SEM (n = 5 biological replicates). F) Representative growth curves (n = 4 biological replicates) of HSP101-kER^TetR-DOZI^ parasites grown at indicated aTc concentrations and treated with DMSO or rapamycin 2 cycles prior. Data are presented as means ± standard deviation from one biological replicate (n = 3 technical replicates). G) Representative EC_50_ assays (n=4 biological replicates) with two-fold dilutions of aTc starting at 500nM. Parasites were treated with DMSO or rapamycin 72 hours prior to adjusting aTc concentrations. Data are presented as means ± standard deviation from one biological replicate (n = 3 technical replicates). EC_50_ graph inset presented as mean ± SEM (n = 4 biological replicates): DMSO=10.58±1.16 nM and rapamycin=9.11±0.29 nM. H) Schematic showing strategy for appending kER and the DDD system to the endogenous C-terminus of HSP101. I) Representative EC_50_ assays (n=4 biological replicates) with two-fold dilutions of TMP starting at 250nM. Parasites were treated with DMSO or rapamycin 72 hours prior to adjusting TMP concentrations. Data are presented as means ± standard deviation from one biological replicate (n = 3 technical replicates). EC_50_ graph inset presented as mean ± SEM (n = 4 biological replicates): DMSO=3.39±0.48 nM and rapamycin 5.48±0.73 nM. Scale bars, 5μm.

If HSP101 functions exclusively in the PV, then knockER-mediated depletion of vacuolar HSP101 levels should hypersensitize parasites to HSP101 knockdown. However, dose-response to aTc was unaltered by rapamycin-treatment (Figure 6G) and the growth defect of KDEL-fused HSP101 parasites maintained at 5 nM aTc was indistinguishable from the DMSO control even over an extended 10-day growth assay (Figure 6F), possibly suggesting that the PV-localized function of HSP101 may not fully account for its role in parasite fitness. To ensure depletion of HSP101 from the vacuole by knockER sensitizes parasites to compromised HSP101 PV function, we generated an HSP101-knockER line where mNG was replaced by a ligand-sensitive DHFR-based destabilization domain (DDD) (Figure 6H) (23). In the absence of trimethoprim (TMP), the DDD blocks HSP101 interaction with EXP2/PTEX150 but not cargo binding, specifically targeting function of the assembled PTEX complex in the PV (23). In contrast to translational knockdown, HSP101-kER^DDD^ parasites were hypersensitized to reduction of TMP levels following rapamycin-treatment, indicating a synergistic effect when the reduction of HSP101 PV levels is combined with disruption of PTEX complex formation (Figure 6I and S7C-E). Taken together, these results illustrate how knockER can be used to study protein function with sub compartmental resolution and may support a distinct HSP101 activity in the ER upstream of its PVM translocation function at the assembled PTEX complex.

## Discussion

Several genetic tools are now available for protein loss-of-function studies in *Plasmodium spp.* (68), including knock-sideways for conditional sequestration of a target protein away from its normal site of function (69, 70). While this system is not currently amenable to parasite proteins that enter the secretory pathway, conditional control of secretory traffic has been achieved in *P. falciparum* by ligand-mediated alteration of the N-terminal transit peptide that controls apicoplast targeting, enabling redirection of plastid proteins into the default secretory pathway (71). Additionally, conditional ER release systems have been developed in model eukaryotes based on ER retrieval machinery (72). In contrast, knockER provides a simple, DiCre-mediated knock-sideways strategy that exploits the ERD2/KDEL receptor (KDELR) system for conditional ER retrieval, providing a generalizable approach for interrogating secreted proteins in genetically tractable eukaryotes.

In this study, we demonstrate knockER simultaneously allows for trafficking and loss-of-function studies. The ability to determine whether a protein traffics from the ER through the *cis* Golgi is an important consideration for nuclear-encoded apicoplast proteins since Golgi-dependent and -independent trafficking routes have been reported (31, 33, 73). While these apicoplast trafficking studies were carried out with reporters and chimeras, knockER allows for monitoring *cis* Golgi transit of endogenous proteins. The sensitivity of ClpB1 to KDEL-mediated ER retrieval demonstrates a Golgi-dependent trafficking route to the apicoplast. Moreover, we show for the first time that this AAA+ chaperone is essential for apicoplast maintenance. ClpB1 is one of four Clp/HSP100 chaperones encoded by *Plasmodium spp*., three of which localize to the parasite plastid while the fourth, HSP101 (previously known as ClpB2) localizes to the PV and ER (43). The other apicoplast-targeted members of this quartet include the nuclear-encoded ClpC, which mediates regulated proteolysis together with the serine protease ClpP (74, 75), and ClpM, now the only uncharacterized member of this group and one of the few proteins encoded on the apicoplast genome. Similar to the well-studied function of its closest orthologs (76), ClpB1 likely acts as a disaggregase that, together with ClpC/P and ClpM, forms a critical proteostasis system required to maintain and regulate the apicoplast proteome (77).

ER perturbations can trigger an integrated stress response leading to cell arrest or death (78, 79). *Plasmodium* possesses one of the widely conserved ER stress sensors, the PERK-eIF2α pathway which halts protein translation and flux through the secretory pathway (20, 80–83). Importantly, conditional ER retrieval of proteins produced from their endogenous locus or by heterologous expression was not generally toxic, enabling loss-of-function studies with knockER. The single notable exception was EXP2, which rapidly killed parasites when retained in the ER independent to loss of its PVM transport function, indicating knockER experiments should be controlled for indirect effects. EXP2-KDEL toxicity was dependent on the amphipathic helix that forms the membrane-spanning region in the oligomeric EXP2 pore (25), suggesting that accumulation of EXP2 in the ER might trigger an unfolded protein response or lead to premature pore formation in the early secretory pathway. More work is required to determine if this outcome is specific to EXP2 but the observation that parasites tolerate SDEL-mediated retention of several TM domain-containing exported proteins indicates this is not a general problem for unnatural ER-retrieval of membrane proteins (84). Indirect fitness defects aside, retrieval of EXP2-KDEL demonstrates that its C-terminus enters the secretory pathway lumen, consistent with a soluble EXP2 trafficking state that precedes membrane integration at the PVM. Regardless of the basis for EXP2-kER toxicity, these experiments highlight that knockER can also be used to investigate membrane topology. While ER-resident type I membrane proteins utilize a different mechanism for retrieval as their C-termini face the cytosol of the cell (85, 86), ERD2/KDELR can retrieve membrane proteins whose C-termini face the lumen (87) and thus knockER should be able to distinguish the topology of single and multi-pass transmembrane domain-containing proteins.

A unique feature of knockER is the ability to redistribute secreted proteins from their terminal destination to the ER. We tagged both HSP101 and the translocon adaptor PTEX150 with knockER, reasoning that ER retrieval of either protein would produce a lethal export defect but that differential impacts on exported cargo trafficking might be exposed. Unexpectedly, ER retrieval of HSP101 had no impact on *P. falciparum* growth or export of representative PEXEL and PNEP proteins *in vitro*. Similar PV depletion of PTEX150 produced the anticipated lethal export block, revealing that HSP101 PV levels are uniquely maintained in excess. As a AAA+ ATPase, HSP101 is the only enzyme in the PTEX core complex and its activity may exceed what is required to support parasite fitness, at least during *in vitro* culture, as previously seen for the aspartic protease Plasmepsin V that licenses proteins for export in the ER (88, 89). Interestingly, parallel experiments in *P. berghei* resulted in greater retention and produced a lethal export defect for both PTEX150 and HSP101, showing knockER is also suitable for conditional mutagenesis in this important model species and suggesting retrieval may be more stringent in *P. berghei*, at least for some proteins. Notably, the strength of ERD2/KDELR binding between different C-terminal retrieval motifs varies by as much as 10-fold in higher eukaryotes (16). While KDEL-mediated retrieval was generally robust across the diverse set of reporters and endogenous proteins surveyed in *P. falciparum* and *P. berghei*, a range of functional variants have been validated or can be inferred in *Plasmodium spp.* (19) and other retrieval motifs may enable tuning the strength of retention by knockER.

Intriguingly, redistribution of HSP101 from the PV to the ER hypersensitized parasites to DDD-mediated HSP101 inactivation but not translational knockdown. While TetR-DOZI knockdown depletes the entire HSP101 pool, HSP101-DDD acts post-translationally by disrupting complex formation with EXP2/PTEX150 but not cargo recognition (23), specifically targeting HSP101 function in PVM translocation. While these results do not enable strong conclusions about potential extra vacuolar activity of HSP101, they suggest that the PV-localized function may not fully account for HSP101’s contribution to parasite fitness, in line with a hypothesized role in exported cargo recognition and/or trafficking at the ER (61). Notably, IBIS1 was not retained in the ER with PbHSP101-KDEL but accumulated in the PV and was not exported into the RBC, indicating that HSP101 is not required to directly chaperone cargo from the ER to the PV. Future work dissecting the specific extra-vacuolar role(s) of HSP101 may resolve the long-standing question of how early events in this pathway that mark proteins for export, such as PEXEL processing during ER entry, are connected to cargo recognition and ultimate PVM translocation into the host cell at the assembled PTEX complex.

## Materials and Methods

### Parasite culture

*P. falciparum* NF54^attB-DiCre^ and derivatives were cultured in RPMI 1640 medium supplemented with 27 mM sodium bicarbonate, 11 mM glucose, 0.37 mM hypoxanthine, 10 µg/ml gentamicin and 0.5% Albumax I (Gibco). Parasites were maintained in deidentified, Institutional Review Board-exempt RBCs obtained from the American National Red Cross. *P. berghei* ANKA clone 2.34 and derivatives were maintained in Swiss Webster mice (Charles River). All experiments involving rodents were reviewed and approved by the Iowa State University Institutional Animal Care and Use Committee.

### Plasmids and genetic modification of *P. falciparum* and *P. berghei*

All cloning was carried out with NEBuilder HIFI DNA assembly (NEB). Primer sequences are given in Table S1. To generate NF54^attB-DiCre^ parasites, a DiCre expression cassette was inserted into the *pfs47* gene in NF54^attB^ parasites (42) using a marker-free CRISPR/Cas9 editing strategy as described (27). Loss of the endogenous *pfs47* locus was monitored with primers P1/2 and P3/4 and integration was confirmed using primers P1/5 and P6/4. To confirm DiCre activity in this line, a loxP-flanked mNG coding sequence with a 3xHA tag was PCR amplified using primers P19/20 and inserted between AvrII and AflII in pLN-ENR-GFP cite (90), resulting in the plasmid JRB416. This plasmid was co-transfected with pINT (90) into NF54^attB-DiCre^ parasites and selection was applied with 2.5 µg/ml blasticidin-S 24 hrs post-transfection.

To generate the kER fusion cassette, homology flanks targeting the 3’ end of *exp2* were amplified from pyPM2GT-EXP2-3xHA-GFP11 (53) using P21/22 and inserted between XhoI and AvrII in pPM2GT-HSP101-3xFLAG (53). The loxPint sequence (28) was amplified with P23/24 and inserted immediately upstream of the 3xFLAG tag at AvrII in this vector. A partial loxPint containing the loxP sequence and 3’ portion of the intron followed by a 3xHA-KDEL tag was then amplified with P25/26 and inserted into EagI, resulting in the plasmid JRB492. Finally, the mNG coding sequencing was amplified from pyPM2GT-EXP2-mNG (91) with P27/28 and inserted into AvrII in JRB492, resulting in plasmid JRB508. For generation of the *exp2* DiCre conditional knockout line, a recoded version of the *exp2* coding sequence with a 3’ 3xHA tag and flanked by loxP sites was amplified using P76/77 from plasmid pEXP2^apt^ (53) and inserted between AvrII/EagI in plasmid JRB492. A 5’ flank immediately upstream of the *exp2* start codon was then amplified from *P. falciparum* gDNA using P78/79 and inserted between AflII/AvrII, replacing the AfllI site with AscI and resulting in the plasmid JRB467. This plasmid was linearized with AscI and co-transfected with pUF-Cas9-EXP2-CT-gRNA into NF54^attB-DiCre^. Selection was applied with 5 nM WR99120 24 hrs post-transfection and clonal lines were isolated by limiting dilution after parasites returned from selection.

To generate the SP-mNG-kER reporter, the *exp2* promoter and signal peptide were amplified from parasite genomic DNA with P29/30 and the mNG-KER cassette was amplified from JRB508 with P31/32 and these amplicons were assembled together in a second PCR reaction with P29/32 and inserted into AscI/ApaI in pKD (41), resulting in the plasmid JRB543. For expression of a second copy of EXP2-kER from the attB locus, the *exp2* promoter and full-length gene were amplified from genomic DNA with P29/33 and inserted between AscI/BamHI in JRB543, resulting in the plasmid JRB526. For expression of a second copy of EXP2-kER lacking the amphipathic helix, a recoded portion of the *exp2* coding sequence beginning at codon 86 (after the amphipathic helix and immediately before the helical body domain) was amplified from pEXP2^apt^ (53) and inserted at BamHI after the EXP2 signal peptide in JRB543, resulting in the plasmid JRB544. These plasmids were each co-transfected with pINT and selected with 2.5 µg/ml blasticidin-S as described above. To relieve translational repression by TetR-DOZI, parasite cultures were supplemented with 500 nM aTc.

For generation of ClpB1-kER and RON6-kER lines, a Cas9 gRNA target was chosen just downstream (*clpB1*, ATTTTTAGAAAAATATTATG) or just upstream (*ron6*, TAATACATACAACGTACCAA) of the stop codon and the gRNA seed sequences were synthesized as primers P34 and P35, respectively, and inserted into the AflII site of pAIO3 (39), resulting in plasmids JRB143 and MAF22, respectively. To target the 3’ end of *clpB1* or *ron6*, a 5’ homology flank (up to but not including the stop codon) was amplified from genomic DNA using P36/37 or P38/39, incorporating synonymous shield mutations in the gRNA target sites. A 3’ homology flank was amplified using P40/41 or P42/43. The corresponding 5’ and 3’ flank amplicons were assembled in a second PCR reaction using P40/37 and P42/39 and inserted between XhoI/AvrII in JRB508, resulting in plasmids MAF20 and MAF27, respectively. These plasmids were linearized with AflII and co-transfected with the corresponding Cas9/gRNA plasmid into NF54^attB-DiCre^. Selection was applied with 5 nM WR99120 24 hrs post-transfection and clonal lines were isolated by limiting dilution after parasites returned from selection.

For generation of PTEX150-kER and HSP101-kER lines, flank assemblies targeting the 3’ end of each gene were amplified using P44/45 from pyPM2GT-PTEX150-3xHA-GFP11 (53) or P46/47 from pyPM2GT-HSP101-3xHA-GFP11 (53) and inserted between XhoI/AvrII in JRB508, resulting in plasmids MAF16 and MAF17, respectively. These plasmids were linearized with AflII and co-transfected with pAIO-PTEX150-CT-gRNA or pAIO-HSP101-CT-gRNA (53), selected with 5 nM WR99120 and cloned after parasites returned. For generation of the EXP2-kER line, plasmid JRB492 (described above) was linearized and co-transfected with pUF-Cas9-EXP2-CT-gRNA (53) and selected with 5 nM WR99120. For generation of HSP101-kER^TetR-DOZI^ lines, flank assembles were amplified using P48/49 and inserted between AscI/BamHI in JRB526 (described above), resulting in plasmid MAF41. This plasmid was linearized with AflII and co-transfected with pAIO-HSP101-CT-gRNA, selected with 2.5 µg/ml blasticidin-S and 500 nM aTc, and cloned after parasites returned. For generation of HSP101-kER^DDD^ lines, the DDD was amplified from plasmid pPM2GT-HSP101-3xFlag-DDD (53) using P70/71 and inserted between AvrII/BsrGI in MAF17, replacing the mNG and resulting in plasmid MAF43. This plasmid was linearized with AflII and co-transfected with pAIO-HSP101-CT-gRNA. Selection was applied with 10 µM TMP 24 hrs post-transfection.

For expression of a second copy of PTEX150-kER from the *attB* locus, the *ptex150* promoter and full-length gene were amplified from genomic DNA with P50/51 and the mNG-KER cassette was amplified from JRB508 with P52/53. These amplicons were assembled together in a second PCR reaction using P50/53 and inserted between ApaI/AflII in pLN-ENR-GFP (90), resulting in the plasmid MAF39. This plasmid was co-transfected with pINT, selected with 2.5 µg/ml blasticidin-S and parasites were cloned after returning.

To generate the line bearing endogenous EXP2-mRuby and HSP101-mNG tags, the plasmid pbEXP2-mRuby3 (92) was first linearized with AflII and co-transfected with pUF-Cas9-EXP2-CT-gRNA (53) into NF54^attB-DiCre^. Selection was applied with 2.5 µg/ml blasticidin-S. Next, the mNG coding sequence was amplified using P54/55 and inserted between AvrII/EagI in pPM2GT-HSP101-3xFLAG (53) to replace the FLAG tag with mNG, resulting in the plasmid JRB325. This plasmid was linearized with AflII and co-transfected with pAIO-HSP101-CT-gRNA into the NF54^attB-DiCre^::EXP2-mRuby parasites. Selection was applied with 5nM WR99120 resulting in the line NF54^attB-DiCre^::EXP2-mRuby+HSP101-mNG.

For generation of PTEX150-kER and HSP101-kER lines in *P. berghei*, the plasmid pBAT was digested with MluI/XhoI and the primer P56 was used as an insert to remove the intervening AvrII, SnaBI and XhoI sites from the MCS. Next, the mRuby3 coding sequence fused to the PbEXP2 signal peptide was amplified with P57/58 and inserted between SwaI/BamHI, replacing the GFP. The mNG-kER tagging sequence was amplified from plasmid JRB508 using P59/60 and inserted between SacII/SphI, replacing the mCherry-3xMYC. Finally, 5’ flanks to tag *P. berghei hsp101* or *ptex150* with mNG-kER were amplified from *P. berghei* genomic DNA using P61/62 or P63/64, respectively, and 3’ flanks were amplified using P65/66 or P67/68, respectively. The corresponding 5’ and 3’ flank amplicons were assembled with an intervening AflII site in a second PCR reaction using P65/62 or P67/64, respectively, and inserted at SacII, resulting in plasmids pPbHSP101-kER+SP-mRuby and pPbPTEX150-kER+SP-mRuby. The plasmids were linearized at AflII and transfected as described (94) into the marker-free HP DiCre line (95). Transfected parasites were reinjected into naïve mice and selection was applied with 0.07 mg/ml pyrimethamine in drinking water provided *ad libitum* 24 hours later. After returning from selection, transfected populations were cloned by limiting dilution IV injection into naïve mice to generate isogenic populations designated PbHSP101-kER and PbPTEX150-kER. To monitor export with an IBIS1-mRuby fusion, the *ibis1* gene and promoter were amplified from *P. berghei* gDNA using P72/73 and the *mruby* cds was amplified using P74/75. These amplicons were assembled in a second PCR reaction using P72/75 and inserted between PvuII/BamHI, resulting in plasmids pPbHSP101-kER+IBIS1-mRuby and pPbPTEX150-kER+IBIS1-mRuby. Plasmids were linearized at AflII and transfected as above.

### P. falciparum growth assays

Growth assays were performed as described previously (22). Briefly, parasites were treated with either DMSO or 10 nM rapamycin for 3 hrs and washed 1X. Then, parasites were seeded between 0.2-1% parasitemia in triplicate and monitored every 48 hrs by flow cytometry on an Attune NxT (ThermoFisher) by nucleic acid staining with PBS containing 0.8 µg/ml acridine orange. For IPP rescue experiments, ClpB1-kER cultures were supplemented with 200 µM IPP. For kER fusions expressed from the *attB* locus, cultures were supplemented with 1 µM aTc 24 hrs prior to rapamycin treatment to relieve TetR-DOZI-aptamer translational repression. For HSP101-kER^TetR-DOZI^ parasites, aTc was removed by washing parasites 7X with 13 mL of media. 500 nM aTc was then supplemented back to control parasites. As required, parasites were sub-cultured to avoid high parasite density, and relative parasitemia at each time point was back-calculated based on actual parasitemia multiplied by the relevant dilution factors. Data were displayed using Prism 9 (GraphPad Software).

For EC_50_ shift assays of HSP101-kER^TetR-DOZI^ and HSP101-kER^DDD^, asynchronous parasites were treated with DMSO or rapamycin before removal of aTc or TMP. After agonist washout as described above, parasites were seeded in triplicate at 0.5% parasitemia with 2-fold dilutions starting from 500 nM aTc or 250 nM TMP and parasite growth was monitored after 72 hrs.

### *P. berghei* kER experiments

Parasite-infected blood was collected by cardiac puncture when parasitemia reached 3-4% and cultured in RPMI supplemented with 20% FBS. Cultures were treated with 10 nM rapamycin or a DMSO vehicle control for 3 hours followed by a media change. After 18 hours of additional culture, parasites were harvested for diagnostic PCR, live fluorescence microscopy and Western blot. Equal numbers of DMSO or rapamycin-treated infected RBCs (∼10,000 parasites per injection) were IV injected into naïve mice and parasitemia was monitored by giemsa-stained blood smear every two days.

For synchronized, *ex vivo* growth assays, parasites were IP injected into mice and allowed to reach 2-3% parasitemia. Parasites gathered from cardiac puncture were then treated with DMSO or 10nM rapamycin 3-4hrs, washed, and allowed to mature into schizonts overnight without further rapamycin treatment. The mature schizonts were magnet purified and tail vein injected into naïve mice treated with phenylhydrazine (1mg/mice) (Sigma, 114715) to allow parasites to egress and invade new RBCs in mice. After 3 hrs to allow parasite egress and re-invasion, synchronous ring-stage parasites were collected by cardiac puncture and allowed to mature overnight *in vitro*. At 24 hrs post injection, parasite development was assessed in Giemsa-stained smears and DNA content was evaluated by Hoechst 34580 staining (1:5,000) on an Attune NxT flow cytometer. To monitor export of IBIS1-mRuby, parasites were IP injected and allowed to grow to 2-3% parasitemia and imaged from tail-blood. Mice were then injected IP with rapamycin (1 mg/kg) and parasites were observed again 24hrs later.

### Microscopy

For live microscopy, *P. falciparum* parasites were treated with either DMSO or 10 nM rapamycin and viewed 24-48 hrs later. Hoechst (33342, Invitrogen; 1:10,000) was used to visualize the nucleus by incubating with parasites 2-5 min prior to visualization. ER-Tracker Red (Invitrogen; 1:1000) was used to visualize the ER by incubating with parasites 30 min at 37°C prior to visualization. For constructs expressed from the *attB* locus under TetR-DOZI-aptamers control, parasites were grown in 500 nM aTc for 24 hrs prior to incubation with DMSO or rapamycin to allow expression. For RON6 parasites, DMSO and rapamycin-treated schizonts were incubated with 50nM ML10 (obtained through BEI resources, NIAID, NIH and supplied by LifeArc, Stevenage, UK: MRT-0207065 (ML10), NR-56525) for 4.5 hrs and visualized by microscopy. For *P. berghei*, synchronized ring-stage parasites were isolated via cardiac puncture, treated with DMSO or rapamycin for 3 hrs and allowed to develop overnight in culture before visualization. For quantification of fluorescence signal, MFI was calculated using the measured Integrated Density value from Fiji (96). The internal MFI was calculated using an ROI within the parasite PV and subtracting that from the whole parasite MFI using an ROI outside the PV boundary. The percent internal fluorescence was then calculated by dividing the internal MFI by the whole parasite MFI.

For protease protection assays on live cells, parasites were treated with 0.03% saponin (Sigma) in 1X PBS for 15 min at 4°C. Purified parasites where then washed once with 5 mL of 1X PBS and once with 5 mL of digestion buffer (50 mM Tris pH 7.5, 150 mM NaCl, 1 mM CaCl_2_) (61). Parasite samples where then split into two identical tubes, one of which was treated with 20 µg/mL of Proteinase K (740506.75, Macherey-Nagel), and incubated at 37°C for 15 min. Proteinase K was removed and samples resuspended with digestion buffer plus protease inhibitor cocktail (Halt 78429, Thermo Scientific) before visualization.

For IFAs of the EXP2- and ClpB1-kER lines, parasites were treated with either DMSO or 10 nM rapamycin and fixed 24 hrs or 144 hrs later as described previously (22). For HSP101-kER^TetR-DOZI^ lines, parasites were synchronized with 5% sorbitol, treated with either DMSO or 10 nM rapamycin, grown with or without 500 nM aTc, and fixed 48 hrs later. Briefly, parasites were fixed with Fixative Solution (4% paraformaldehyde, 0.0075% glutaraldehyde in PBS) for 25 min at room temperature in the dark. Parasites were then permeabilized with 0.1% Triton in PBS for 10 minutes at room temperature while rocking. Parasites were washed 1X with PBS and then blocked with 3% BSA in PBS for 1hr at room temperature. Primary antibody was applied in 3% BSA solution for 1 hr at room temperature, washed 2X with PBS, followed by secondary antibody incubation for 1 hr at room temperature. After 2X washes with PBS, parasites were adhered onto Poly-L-lysine coated coverslips and then mounted onto slides. For parasites probed with SBP1, parasites were acetone fixed as in (39). Briefly, thin smears were immersed in 100% acetone for 2 min. Smears were washed 1X with PBS and then blocked with 3% BSA in PBS for 1 hr at room temperature. Primary antibody was applied in 3% BSA solution for 1 hr at room temperature, washed 2X with PBS, followed by secondary antibody incubation for 1 hr at room temperature. After 2X washes with PBS, slides were mounted with DAPI, covered with a coverslip and sealed. The antibodies used for IFA were: mouse anti-HA antibody (clone 16B12, BioLegend; 1:200), rabbit polyclonal anti-HA SG77 (ThermoFisher; 1:500), rabbit anti-PfGRP78 (BiP) (MRA-1246, BEI Resources, NIAID, NIH; 1:100), mouse monoclonal anti-FLAG clone M2 (Sigma-Aldrich; 1:200), rabbit monoclonal anti-Flag 8H8L17 (ThermoFisher; 1:500), mouse anti-HRP2 clone 2G12 (1:500) (97), rabbit polyclonal anti-SBP1 BR28 (1:500) (98) and rabbit polyclonal anti-ACP1 (1:500 IFA) (37, 99). Secondary antibodies used were anti-mouse antibody conjugated to Alexa Fluor 488 or 546 and anti-rabbit antibody conjugated to Alexa Fluor 488, (Life Technologies; 1:1000). Parasites were mounted with ProLong diamond with 4’,6’-diamidino-2-phenylindole (DAPI) (P36931; Invitrogen) and imaged on an Axio Observer 7 equipped with an Axiocam 702 mono camera and Zen 2.6 Pro software (Zeiss) using the same exposure times for all images across sample groups and experimental replicates. Image processing, analysis, and display were performed using Zeiss Blue software and Adobe Photoshop. Equivalent adjustments to brightness and contrast were made within sample groups for display purposes.

### Quantification of protein knockdown

To evaluate the level of HSP101 protein knockdown, HSP101-kER^TetR-DOZI^ parasites were synchronized via magnet purification and treated with DMSO or rapamycin before removal of aTc. After aTc removal, parasites were seeded at ∼1% parasitemia with different dilutions of aTc and allowed to grow for 48 hrs. Using an Attune NxT flow cytometer, infected RBCs were gated using Hoechst 34580 (Invitrogen; 1:5,000) and mNG fluorescence of infected RBCs was concurrently measured. The same procedure was applied for saponin-released parasites after protease treatment as described above.

### Western blotting

Western blotting was performed as described previously (39, 53). Briefly, parasite-infected RBCs were selectively permeabilized by treatment with ice-cold 0.03% saponin in PBS for 15 min followed by lysis with RIPA and centrifugation to remove hemozoin. For constructs expressed from the *attB* locus under control of TetR-DOZI-aptamers, parasites were grown in 500 nM aTc for 24 hrs prior to incubation with DMSO or rapamycin to allow for expression. For the HSP101-kER^TetR-DOZI^ lines, parasites were treated with DMSO or rapamycin before removal of aTc. After aTc removal, parasites were seeded at ∼1% parasitemia with different dilutions of aTc and allowed to grow for 48 hrs before processing. The antibodies used in this study were mouse anti-HA antibody (clone 16B12, BioLegend; 1:500), rabbit polyclonal anti-HA SG77 (ThermoFisher; 1:500), mouse anti-FLAG (Clone M2, Sigma-Aldrich; 1:500), rabbit polyclonal anti-*Plasmodium* Aldolase ab207494 (Abcam; 1:500) and mouse monoclonal anti-EXP2 clone 7.7 (1:500) (100). The secondary antibodies used were IRDye 680CW goat anti-rabbit IgG and IRDye 800CW goat anti-mouse IgG (Li-COR Biosciences; 1:20,000). Western blot images were acquired using an Odyssey Clx infrared imaging system and processed with Image Studio software (Li-COR Biosciences). Full length versions of cropped blots can be seen in Figure S8.

## Acknowledgements

This work was supported by NIH grant HL133453 to JRB. The funders had no role in study design, data collection and interpretation, or the decision to submit the work for publication. We thank J. McBride, D. Cavanaugh and EMRR for the EXP2 antibody and A. Waters for the HP DiCre *P. berghei* parasites.

## Author contributions

M.A.F. and J.R.B designed research; M.A.F., T.H., L.J.C. and J.R.B performed research; M.A.F. and J.R.B analyzed data; M.A.F. and J.R.B. wrote the paper.

