## Supplemental Figures for "Knock-sideways by inducible ER retrieval enables a novel approach for studying *Plasmodium* secreted proteins"

**Supplemental Figures and Tables for Fierro *et al***

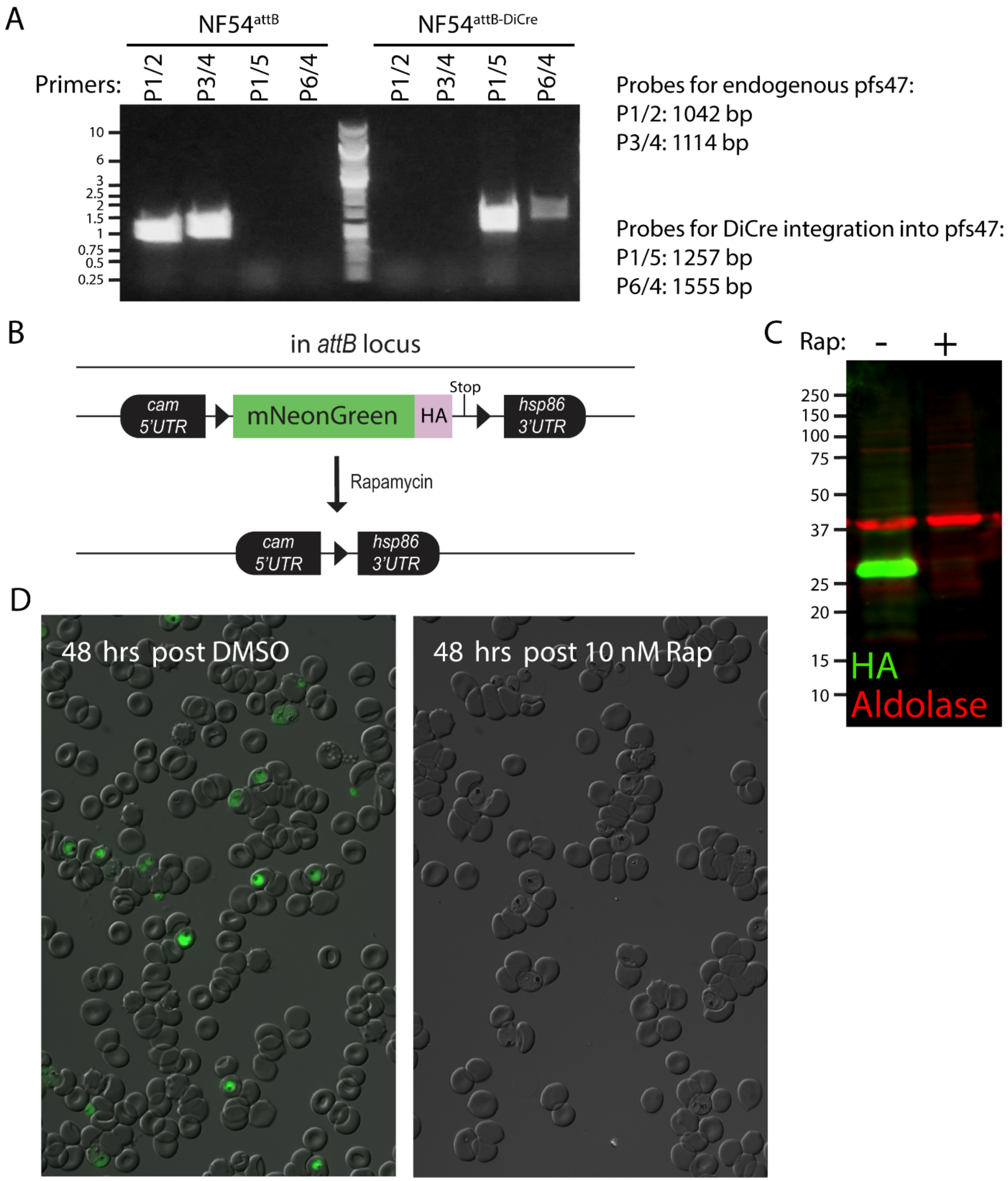


**Figure S1. Generation and validation of NF54$^{attB\text{-}DiCre}$ parasites.** A) Diagnostic PCR showing DiCre cassette integration into the *pfs47* locus using primers P1/2 and P3/4 to detect unmodified locus, and primers P1/5 and P6/4 to detect integration. B) Schematic of reporter integrated at *attB*

in *cg6* to test DiCre functionality in $NF54^{attB\text{-}DiCre}$. C) Western blot and D) live microscopy of parasites 48hrs after 3hr treatment with DMSO or 10nM rapamycin. Molecular weights are predicted to be 30 kDa for mNG-3xHA and 40.1 kDa for aldolase.

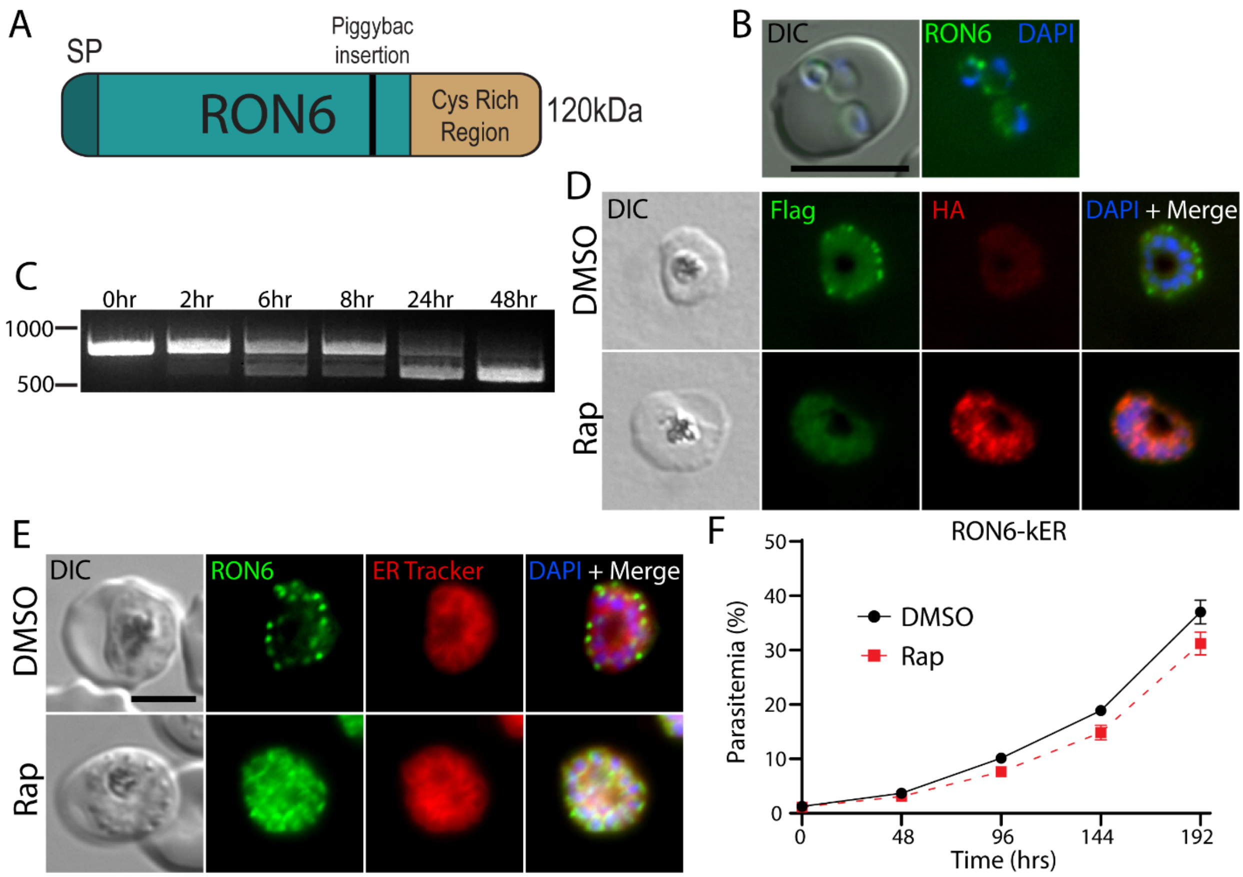


**Figure S2. Inducible ER retention of rhoptry protein RON6.** A) Schematic of rhoptry neck protein RON6. SP= signal peptide; black bar= site of piggyBac insertion in RON6 (46). B) Live microscopy of ring-stage RON6-kER parasites showing peripheral RON6-mNG signal, indicating RON6 localization to the PV after invasion. C) Time course of excision following rapamycin treatment detected by PCR using primers P7/8. D) Representative IFA of a RON6-kER schizont showing conversion of Flag to HA and retention of the HA-tagged species in the ER following rapamycin treatment. Synchronized parasites were treated with DMSO or rapamycin at 24 hpi and allowed to develop for 72 hrs to second cycle schizonts before fixation. At least 40 iRBCs were assessed across multiple fields in each condition; all rapamycin-treated parasites showed conversion to the HA-KDEL tag. E) Live microscopy of DMSO or rapamycin-treated RON6-kER parasites 48hrs post-treatment, treated with 25nM ML10 for 4hrs (BEI Resources) to prevent egress and arrest parasites as terminal schizonts. F) Representative growth of asynchronous RON6-kER parasites (n=3 biological replicates) treated with DMSO or rapamycin. Data are presented as means ± standard deviation from one biological replicate (n = 3 technical replicates). Scale bar, 5μm.

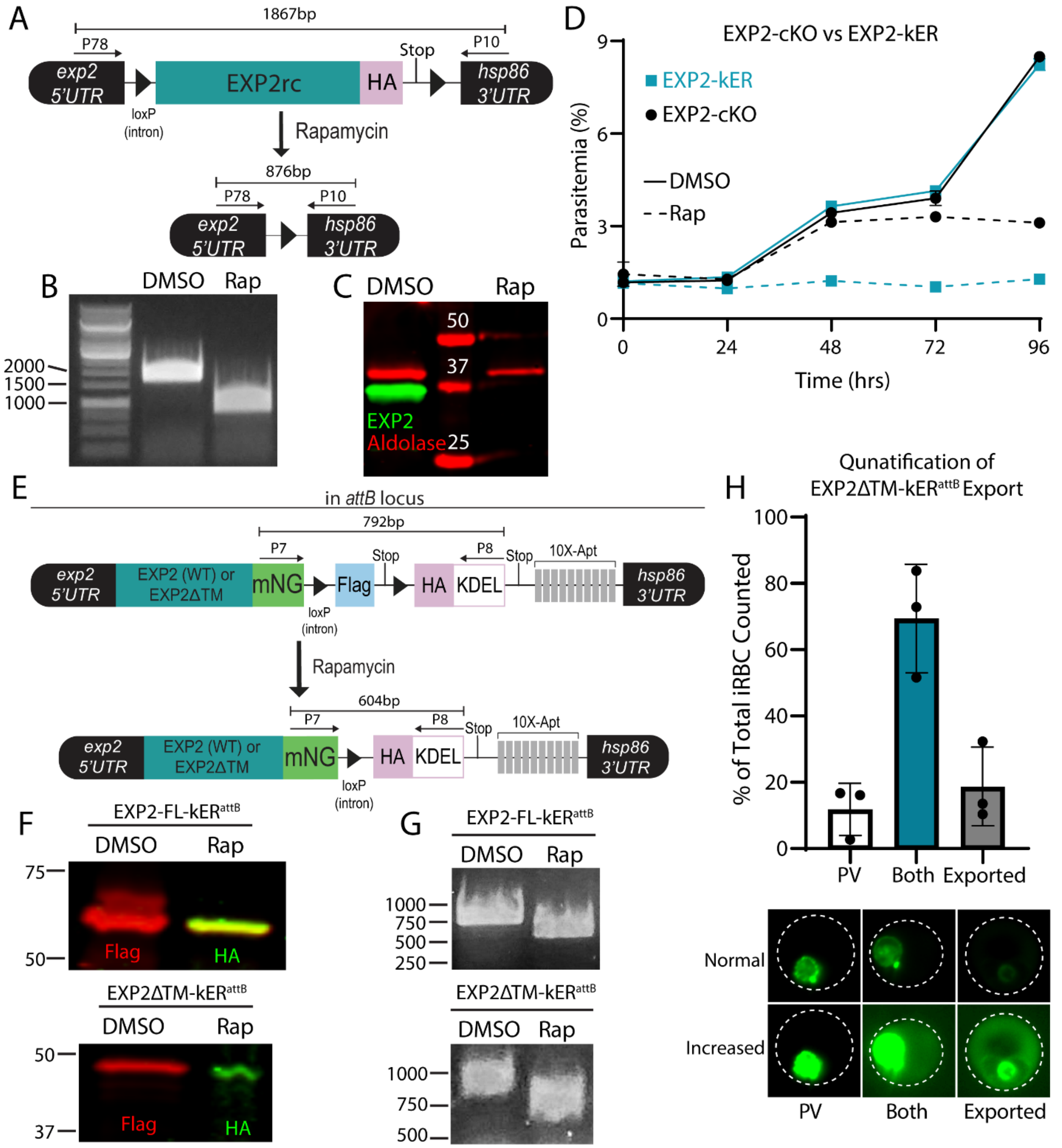


**Figure S3. Supporting data for Figure 3.** A) Schematic showing the modified, floxed endogenous *exp2* locus for DiCre-based conditional knockout in EXP2-cKO parasites. EXP2rc; recoded EXP2 sequence. B) PCR showing excision of the *exp2* gene in EXP2-cKO parasites 48 hours after rapamycin treatment using primers P78/P10. C) Western blot 48 hours after treatment of EXP2-cKO parasites with DMSO or rapamycin showing loss of EXP2 expression. Aldolase serves as a loading control. Molecular weights are predicted to be 40.1 kDa for aldolase and 33.8 kDa for EXP2-3xHA after signal peptide cleavage. D) Representative comparative growth assay of EXP2-kER and EXP2-cKO parasites (n=2 biological replicates). Synchronized, ring-stage parasites of each line were treated with DMSO or rapamycin at time 0. Data are presented as means ± standard deviation from one biological replicate (n = 3 technical replicates). E) Schematic showing cassette for second copy expression of full-length EXP2 or a version lacking the amphipathic helix with kER fusion from the *attB* site of chromosome 6. F) Western blots from second copy EXP2-kER parasites maintained in media supplemented with 500 nM aTc. Samples were taken 24hrs after DMSO or rapamycin treatment showing conversion from Flag to HA. Molecular weights after signal peptide cleavage are predicted to be 61.4 kDa for EXP2-FL-3xFLAG, 61.5 kDa for EXP2-FL-3xHA-KDEL, 54.4 kDa for EXP2ΔTM-3xFLAG and 54.5 kDa for EXP2ΔTM-3xHA-KDEL**.** G) PCR showing excision in EXP2-FL-kER$^{attB}$ and EXP2ΔTM-kER$^{attB}$ parasites 24hrs after rapamycin treatment using primers P7/8. H) Quantification of EXP2ΔTM-mNG export in EXP2ΔTM-kER$^{attB}$ parasites without rapamycin treatment (n=3 biological replicates). Infected RBCs (iRBCs) were scored as having strict PV retention without apparent export (PV), both exported and PV signal (both), or full export without apparent PV signal (full). Error bars indicate mean ± standard deviation. Representative images are shown below and displayed with normal or increased brightness and contrast to clearly show the fractions of mNG signal localized to the PV, digestive vacuole and RBC cytosol.

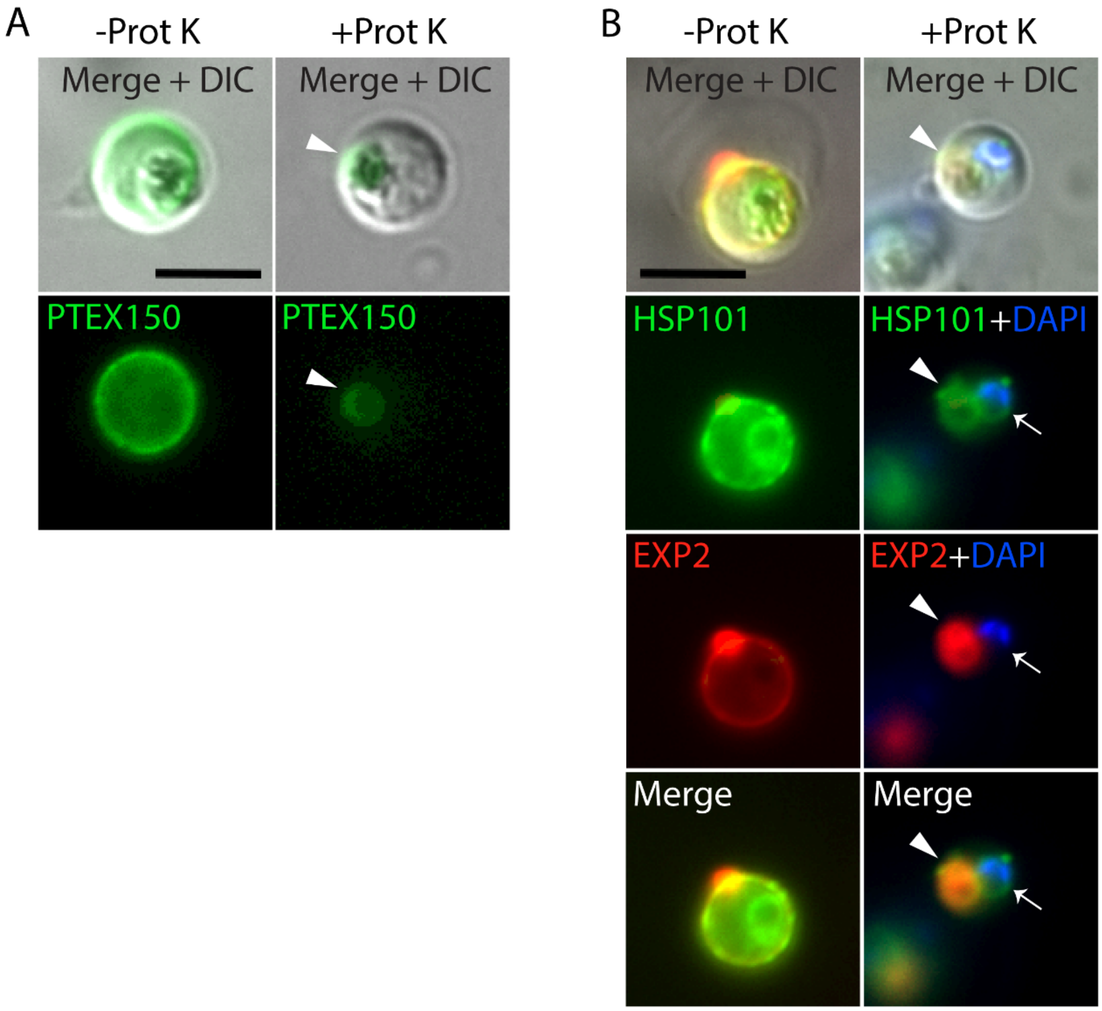


**Figure S4. Protease protection assay on live cells verifies an internal, perinuclear pool of HSP101 but not EXP2 or PTEX150.** Live microscopy of parasite lines with endogenous fluorescent protein fusions A) PTEX150-mNG-3xFLAG or B) EXP2-mRuby3 and HSP101-mNG treated with saponin +/- Proteinase K. Images are representative of n=2 biological replicates. Arrowhead: protease-protected fluorescence within the digestive vacuole from endocytosis typically seen for PV proteins; arrow: protease-protected perinuclear fluorescence unique to HSP101. Scale bar, 5µm.

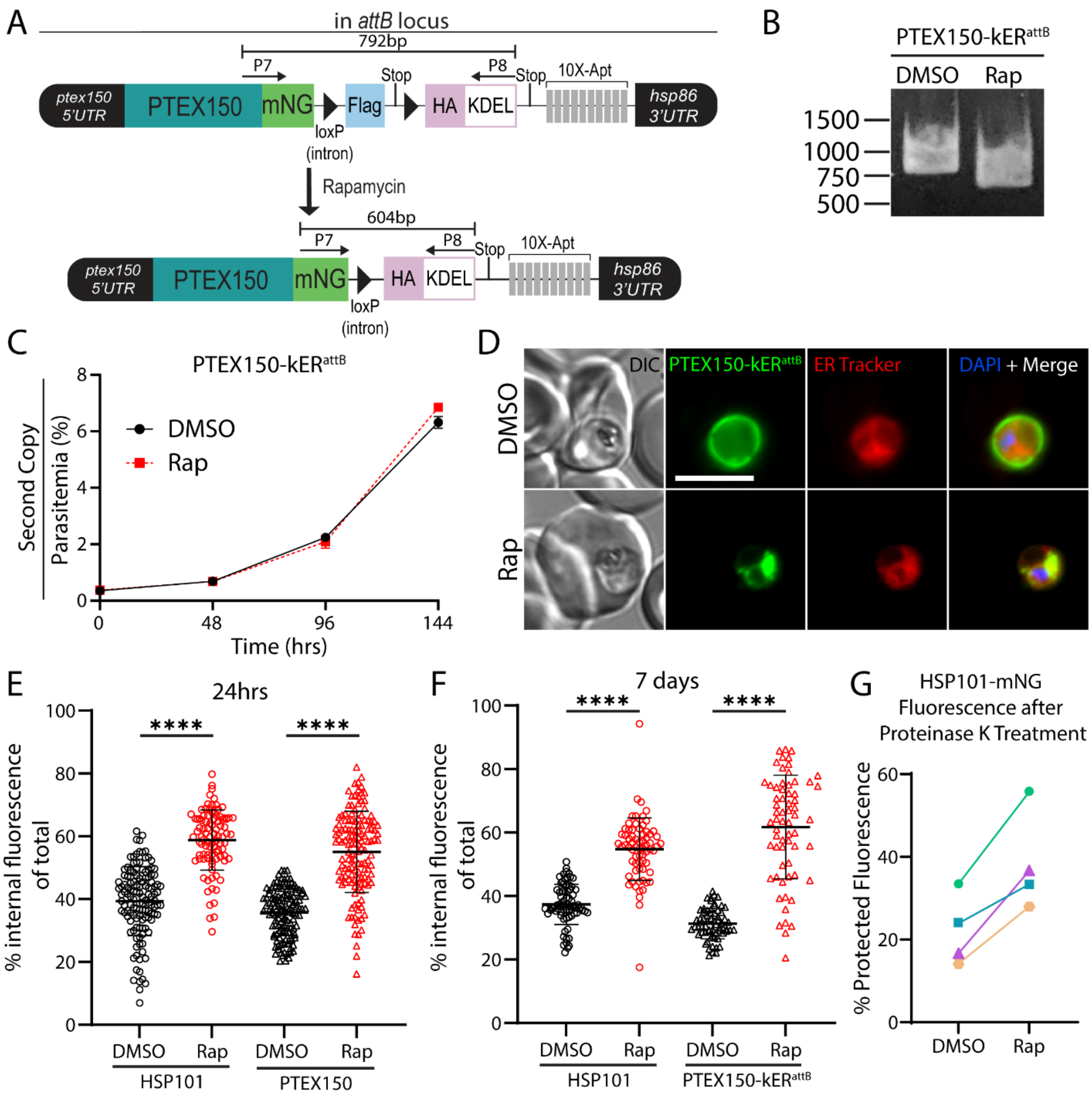


**Figure S5. Supporting data for Figure 4.** A) Schematic showing cassette for second copy expression of PTEX150-kER from the *attB* site of chromosome 6. B) PCR showing excision in PTEX150-kER$^{attB}$ parasites 24hrs after rapamycin treatment using primers P7/8. C) Representative growth curves of asynchronous, second copy PTEX150-kER parasites (n=3 biological replicates) treated with DMSO or rapamycin. Data are presented as means ± standard deviation from one biological replicate (n = 3 technical replicates). D) Live microscopy of second copy PTEX150-kER parasites 24hrs post-treatment with DMSO or rapamycin. PTEX150-kER$^{attB}$

cultures were maintained in media supplemented with 500 nM aTc. E) Quantification of percent internal mNG fluorescence in live microcopy images of endogenously tagged HSP101- and PTEX150-kER parasites 24hrs post-treatment with DMSO or rapamycin. Data are pooled from 3 (HSP101) or 2 (PTEX150) independent experiments and bars indicate mean ± standard deviation (****, $P<0.0001$; unpaired t test). F) Quantification of percent internal mNG fluorescence in live microcopy images of endogenously tagged HSP101-kER and second copy PTEX150-kER parasites lines 168hrs post-treatment with DMSO or rapamycin. Data are pooled from 2 independent experiments and bars indicate mean ± standard deviation (****, $P<0.0001$; unpaired t test). G) Mean fluorescence intensity quantification from proteinase K treated parasites. The amount of protected fluorescent signal (percent of total before protease treatment) was determined by dividing the MFI after proteinase K treatment by the MFI before proteinase K treatment. Colors represent different replicates.

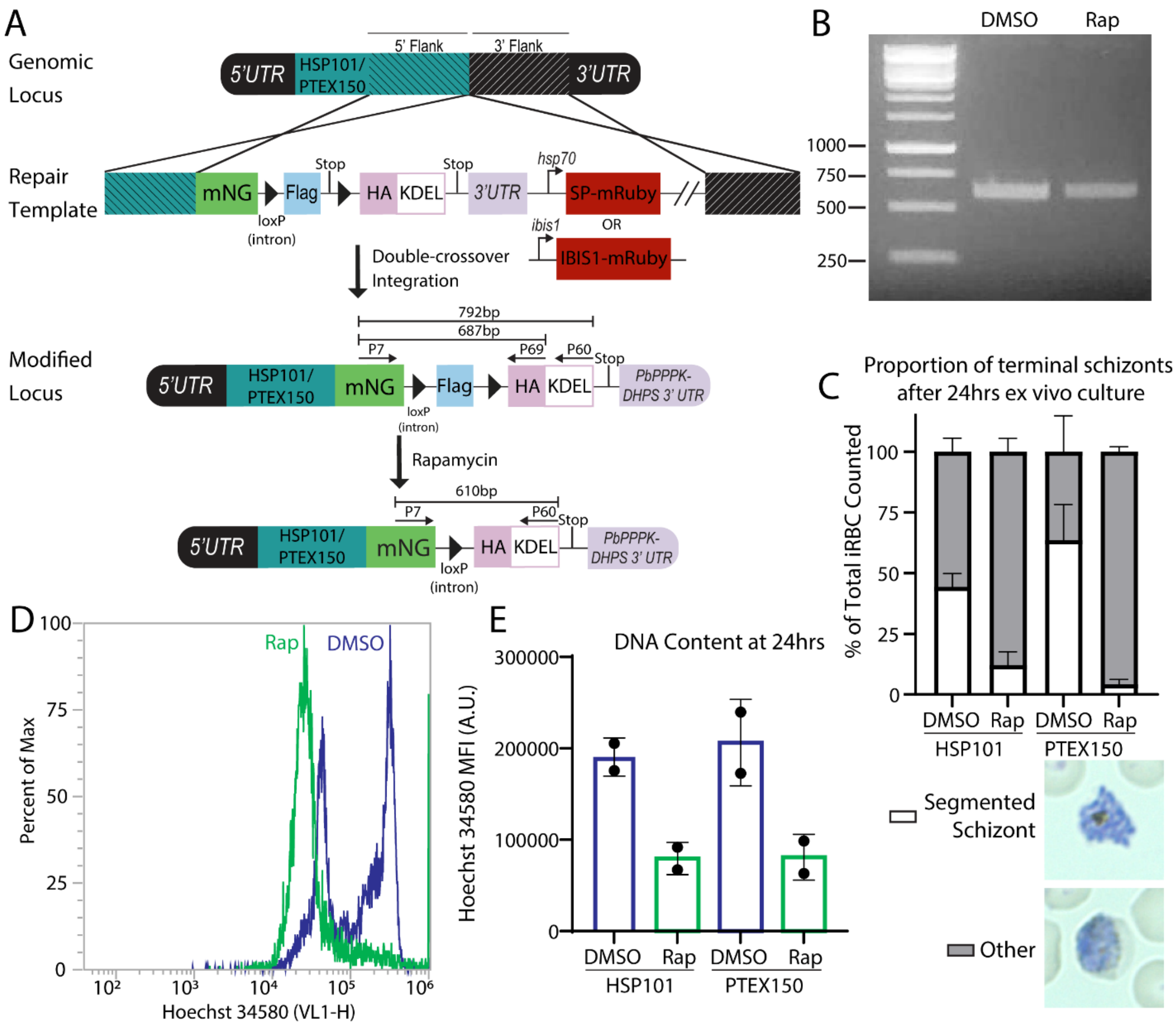


**Figure S6. Supporting data for Figure 5.** A) Schematic showing strategy for appending kER to the endogenous C-terminus of *P. berghei hsp101* and *ptex150*. Tagging plasmids include a downstream cassette for expression of mRuby fused to the EXP2 signal peptide for default secretion into the PV or IBIS1-mRuby, which is exported into the RBC. B) PCR using P7/P69 to evaluate excision of the kER cassette in parasites taken from mice injected with DMSO-treated PbHSP101-kER (collected on day 8) or rapamycin-treated PbHSP101-kER (taken from the single mouse where parasites appeared on day 10). The results indicate that the kER cassette was unexcised in the parasites observed in the mouse that became patent on day 10. C-E) Assessment of PbHSP101-kER and PbPTEX150-kER parasite development in *ex vivo* culture. Synchronous ring-stage parasites (≤3 hpi) treated with DMSO or rapamycin in the preceding cycle were allowed to develop *ex vivo* for 24 hrs. Parasite development was assessed by C) the proportion of terminal, segmented schizonts from Giemsa-stained smears (representative images shown below) and D,E) flow cytometry analysis of parasite DNA content. For flow cytometry analysis, 24 hr *ex vivo* cultures were stained with Hoechst 34580 and infected RBCs were gated using mNG fluorescence and Hoechst 34580 fluorescence (excited with a 405 nm laser) was collected from this population. D) Representative histogram of Hoechst 34580 fluorescence from one PbPTEX150-kER experiment. E) DNA content of rapamycin-treated parasites is substantially decreased compared to DMSO controls, in agreement with failure to normally complete schizogony and consistent with a major developmental defect in PbHSP101-KDEL and PbPTEX150-KDEL during *ex vivo* culture. Points represent MFI from 5,000 iRBCs in each of two independent experiments. Error bars indicate standard deviation.

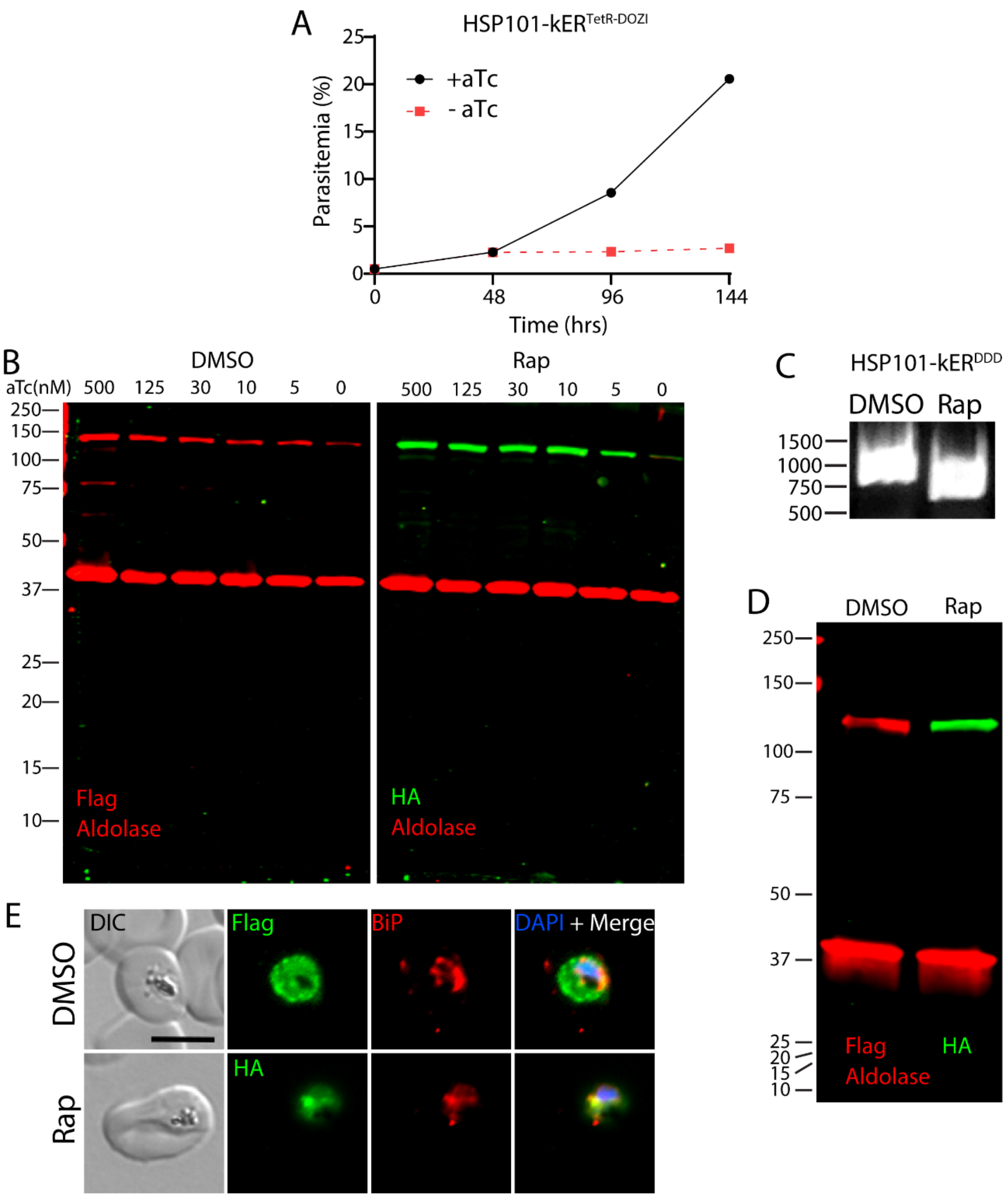


**Figure S7. Supporting data for Figure 6.** A) Representative growth curve (n = 3 biological replicates) of HSP101-kER$^{TetR-DOZI}$ parasites grown with or without aTc. Data are presented as means ± standard deviation of one biological replicate (n = 3 technical replicates). B) Western blot of HSP101-kER$^{TetR-DOZI}$ parasites grown 48hrs at indicated aTc concentrations. Parasites grown in 500 nM aTc were treated with DMSO or rapamycin 24hrs before aTc was washed out and restored at indicated concentrations. Results are representative of 4 independent experiments.

Molecular weights after signal peptide cleavage are predicted to be 130.5 kDa for HSP101-mNG-3xFLAG and 130.6 kDa for HSP101-mNG-3xHA-KDEL. C) PCR showing excision in HSP101-kER$^{DDD}$ parasites 72hrs after rapamycin treatment using primers P70/8. D) Western blot of HSP101-kER$^{DDD}$ parasites 2.5 cycles post-treatment with DMSO or rapamycin. Molecular weights after signal peptide cleavage are predicted to be 122.1 kDa for HSP101-DDD-3xFLAG, 122.2 kDa for HSP101-DDD-3xHA-KDEL and 40.1 kDa for aldolase. E) IFA of HSP101-kER$^{DDD}$ parasites 2.5 cycles post-treatment with DMSO or rapamycin. BiP serves as a marker for the ER.

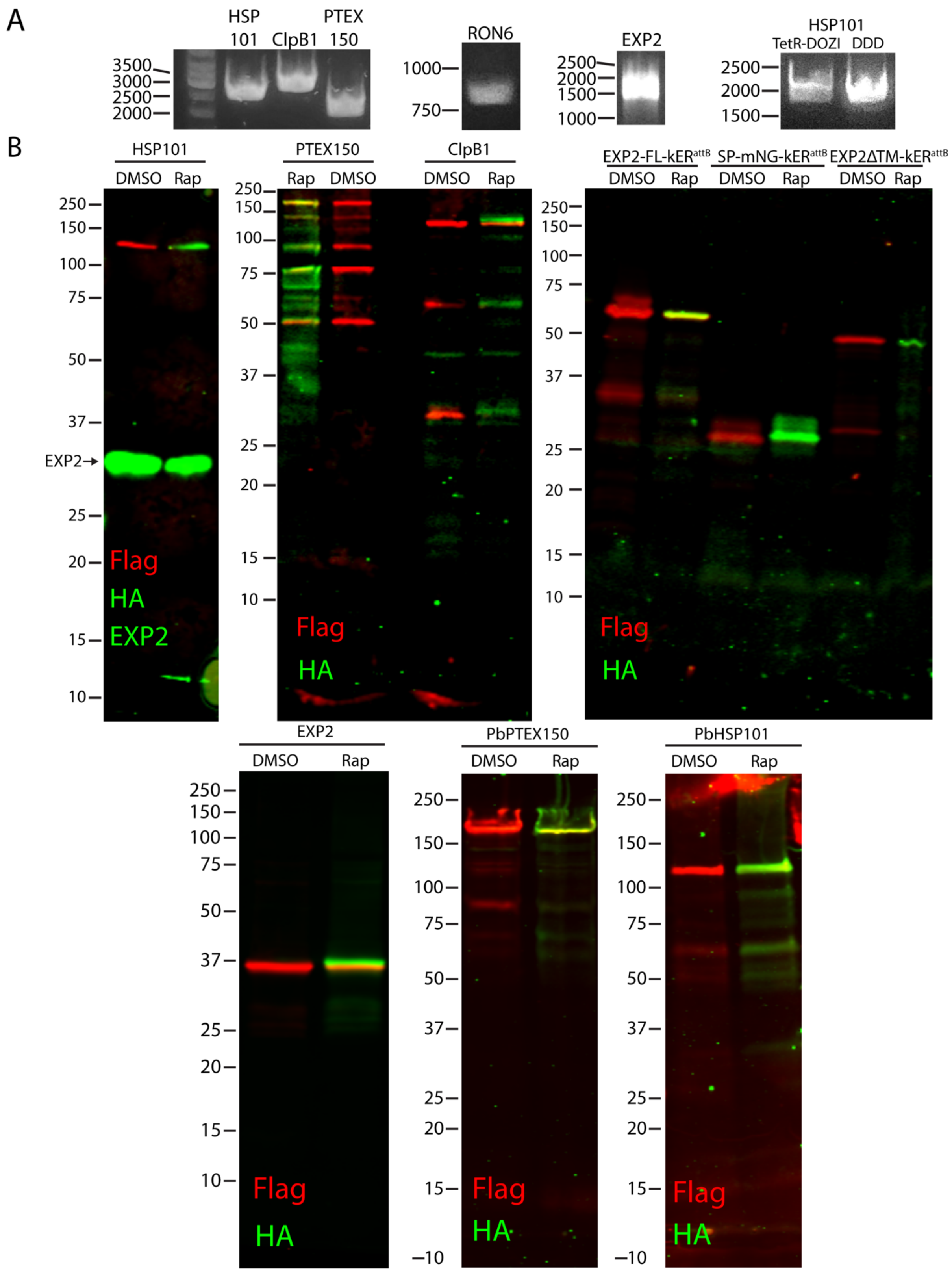


**Figure S8. Integration PCRs and Uncropped Western blots** A) Integration PCR for endogenous kER tags using primers P11/P12 (HSP101), P13/P12 (ClpB1), P14/P12 (PTEX150), P15/P16 (RON6), P17/P8 (EXP2), P11/P12 ($HSP101^{TetR-DOZI}$), and P11/P71 ($HSP101^{DDD}$). B) Uncropped Western blots.

**Table S1.** Sequences of primers used in this study.

| Name | Sequence |
|---|---|
| P1 | ATTGCATACACATAAATATTTGTGTTGTAC |
| P2 | CACATACGTATTGTGTTGAGCTTAATAG |
| P3 | CTAATGTTAAGCCAACTGTAGTTGGG |
| P4 | GATGCGATATGTAATTCCATTACTGC |
| P5 | CCATTTACACATAAATGTCACACAAAAGAG |
| P6 | GAAATAATTTCATATACACACATATAC |
| P7 | GCTACACCTACGAGGGAAGC |
| P8 | ATATATTAATAACTCGACGCGGCCGTTATAATTCGTCCTTGGCATAATCTGGAACATCGTAAGGATACG |
| P9 | CAGATGACGACGATGATGATGACG |
| P10 | GTATATTGGGGTGATGATAAAATGAAAG |
| P11 | CGAAAACTTTTATGGTATTAATATAACAG |
| P12 | GTAAGTCTTCTTCGACCTGCAC |
| P13 | GATGAAATCCATACTGTTGTGGGAGC |
| P14 | CGATGATGATGATGATTTGGATGAAG |
| P15 | CTTGACATGAATGAAACCATACAAAAATTAAAATCCGATAGTG |
| P16 | GACGATGTACATTTTGTACCTTCCCAATTTGTTACTATAAAAGTAAGACATC |
| P17 | CACTCCTCCACTTCCCCTAGGTCCCGTTTTTTGTTTCTCGTGCAAGGGTAATACATACAACGTACCAATTGTGTC |
| P18 | CAGATGACGACGATGATGATGACG |
| P19 | ACCTAATAGAAATATATCACCTAGGATAACTTCGTATAGCATACATTATACGAAGTTATATGGTGAGCAAGGGCGAGGAG |
| P20 | CCTTACGATGTTCCAGATTATGCCTGAATAACTTCGTATAGCATACATTATACGAAGTTATCTTAAGGTCGAGTTATATAATATAT |
| P21 | CACTATAGAACTCGAGGGAGAAACAATCTTTTATATAAAATGTACAGAGTTTGAAAG |
| P22 | GTGGAGATTTATTTAAAAATGAAAAAAAAGATGAAAATAAAGAACCTAGGGACTACAAGGACGACGACG |
| P23 | AAAAAGATGAAAATAAAGAACCTAGGGTAAATAAAAAAAATAATATACAATAACTTCGTATAGCATACATTATACGAAG |
| P24 | TCGTCGTCGTCCTTGTAGTCGCTAGCGTTGCCGTTATTCTGGCTAGCC |
| P25 | AAGATGATGATGATAAATGAACTAGTATAACTTCGTATAGCATACATTATACGAAGTTATTATATATG |
| P26 | CGTATCCTTACGATGTTCCAGATTATGCCAAGGACGAATTATAACGGCCGCGTCGAGTTATTAATATAT |
| P27 | GATGAAAATAAAGAACCTAGGGGAAGTGGAGGAGTGAGCAAGGGCGAGGAGGATAAC |
| P28 | ATATTATTTTTTTTATTTACCTTGTACAGCTCGTCCATGCCCATC |
| P29 | TCCAATGGCCCCTTTCCGGGCGCGCCCCTTGTAAATTTTAATAGTTATAATATAATATTATTTACTTCTACATCCACTG |
| P30 | CGCCCTTGCTCACTCCTCCGGATCCAACTACAGTGTTTGTATTTTTATATACGAAGAATAACAAAAAAAAGG |
| P31 | GGATCCGGAGGAGTGAGCAAGGGCG |
| P32 | CTCTGCCTTGCCAGACAGTGGGCCCTTATAATTCGTCCTTGGCATAATCTGGAACATCG |
| P33 | CGCCCTTGCTCACTCCTCCGGATCCTTCTTTATTTTCATCTTTTTTTTCATTTTTAAATAAATCTCCACTGGC |
| P34 | CATATTAAGTATATAATATTATTTTTAGAAAAATATTATGGTTTTAGAGCTAGAAATAGC |
| P35 | CATATTAAGTATATAATATTTAATACATACAACGTACCAAGTTTTAGAGCTAGAAATAGC |
| P36 | GTAACCTTGTGCATATCAGAAAATTGTTTGCACTCCTTAAGGATGAAGTTACAAGTGAAGATATTGTTAATATTGTAAGTATGTCTACC |
| P37 | CACTCCTCCACTTCCCCTAGGACTTTTTGAAAAGTGCAACTTCTGATCTTTCAGAG |
| P38 | AGCAATACATACATAATACCTTAAGGGATATAATTTACATGAATGTTTACAATTTTTGGTCGTC |
| P39 | CACTCCTCCACTTCCCCTAGGTCCCGTTTTTTGTTTCTCGTGCAAGGGTAATACATACAACGTACCAATTGTGTC |
| P40 | CACTATAGAACTCGAGCAAGTACATAAAGAAAATGCTTATCAATCATCGATTTA |
| P41 | GGTAGACATACTTACAATATTAACAATATCTTCACTTGTAACTTCATCCTTAAGGAGTGCAAACAATTTTCTGATATGCACAAGGTTAC |
| P42 | TTTAGGTGACACTATAGAACTCGAGCAATATGTATATATATATATATGCAAATGTATAAATCTACATATGCATATTGTC |
| P43 | ATGTAAATTATATCCCTTAAGGTATTATGTATGTATTGCTTAATTTAATAATGATTATCCATTTTTTTAATTTTTG |
| P44 | TTTAGGTGACACTATAGAACTCGAGGGTATAGAAAAAATATATAATATTTATATGCTTTTCTGCCAAATTTGC |
| P45 | CACTCCTCCACTTCCCCTAGGGTTATCATCTTCTTCTTCGTCTAATTCTTCTTCATCATTAGAATCATCG |
| P46 | GGTGACACTATAGAACTCGAGAATTACGCATATATATATATATATATATATATAACATGGGTTGTATAATTTAAAAG |
| P47 | CACTCCTCCACTTCCCCTAGGGGTCTTAGATAAGTTTATAACTAAGTTTTTAGCTTTAC |
| P48 | TCCAATGGCCCCTTTCCGGGCGCGCC |
| P49 | CGCCCTTGCTCACTCCTCCGGATCCGGTCTTAGATAAGTTTATAACTAAGTTTTTAGCTTTACTATTATAATCAAC |
| P50 | ATTAGCTAAGCATGCGGGCCCGTTGTTTTTCTCTTTGTGGTCAAAATAAGTAAAATTTTATAAATTC |
| P51 | CTCGCCCTTGCTCACCTCGAGATTGTCGTCCTCTTCTTCGTCCAATTCTTCTTCATC |
| P52 | CGACAATCTCGAGGTGAGCAAGGGCGAGGAGG |
| P53 | CCTTACGATGTTCCAGATTATGCCAAGGACGAATTATAACTTAAGGTCGAGTTATATAAT |
| P54 | TATAAACTTATCTAAGACCCCTAGGGGAAGTGGAGGAGTGAGCAAGG |
| P55 | GGCATGGACGAGCTGTACAAGTGACGGCCGCGTCGAGTTATATAATATA |
| P56 | AATATATAAGTAAGAAAAAACGCGTCCCGGGAAGCTTATCGATGG |
| P57 | AATTACAATTACAATTATTTAAATATGAAGATTCGTTATTTTTTGTCTCTTTTTTTATTTTATGCGACCTATAAGCATACAACAGCTAGCCCATGGGGAAGTGGAGGAGTGTCTAAGGGC |
| P58 | CTTGCACACCTTTTAGCTAGGATCCTTAGGCATAATCTGGAACATCGTAAGGATACG |
| P59 | TACTTCTCGCGAGCGCGCCCGCGGGGAAGTGGAGGAGTGAGCAAGGG |
| P60 | CGATGTTCCAGATTATGCCAAGGACGAATTATAATCTAGAACTTATTGAAGAAAAAAAT |
| P61 | GCATTAGGATTAGATATTTGCGGTTTGGTACAACATATTAGCTTAAGCTCATAATGCTATGAATGTAGATTCACTTTATCAAGAGC |
| P62 | TGCTCACTCCTCCACTTCCCCTAGGTGACAATGAAAGATTAATAACAATGTTGTTGATTGTTTTGTTGTAATCAACATATACATCCATATCATC |
| P63 | GTAGCATGGTGTGATTTATATCTCCTCATTTGTTGAACCTTAAGCAATTAATTCAAGACAATATACTCGATTATAGTGATTTAACAG |
| P64 | TGCTCACTCCTCCACTTCCCCTAGGTTCATCTTCATCTTCATCTTCTGGGTAATCTTCTAATAATTCATTATCACTGTTGCC |
| P65 | TACTTCTCGCGAGCGCGCCCGCGGCAAACACGATATGTTGCATGTATATATATTTATCTTTTATTATTTGCTTATCG |
| P66 | GCTCTTGATAAAGTGAATCTACATTCATAGCATTATGAGCTTAAGCTAATATGTTGTACCAAACCGCAAATATCTAATCCTAATGC |
| P67 | TACTTCTCGCGAGCGCGCCCGCGGGAATATTGCTTTCCCTATATATAATTATAAAATTGAAAAAAGAGAAGCTGAC |
| P68 | CTGTTAAATCACTATAATCGAGTATATTGTCTTGAATTAATTGCTTAAGGTTCAACAAATGAGGAGATATAAATCACACCATGCTAC |
| P69 | CCGGGACGTCGTACGGG |
| P70 | TATAAACTTATCTAAGACCCCTAGGGGAAGTGGAGGAATCAGTCTGATTGCGGCGTTAGCGGTAGATC |
| P71 | ATTTTTTTTATTTACTGTACATCGCCGCTCCAGAATCTCAAAGCAATAGCTGTGAGAGTTC |
| P72 | TTTGTTGTGCTAAGCACCAGCTGGAATGTCGATGTTTTATAAGGGCATAAAAATGTAG |
| P73 | TTTCCTTGATCAGCTCTTCGCCCTTAGACACTCCTCCACTTCCGGCGCCCATAGGTTTTGCTCTACAAAATATGTTAGATTTATTCAAATTTTTTCTTTC |
| P74 | GGAAGTGGAGGAGTGTCTAAGGGC |
| P75 | CTTGCACACCTTTTAGCTAGGATCCTTACTTGTACAGCTCGTCCATGCCAC |
| P76 | AAAGATGAAAATAAAGAACCTAGGATAACTTCGTATAGCATACATTATACGAAGTTATGCTAGCATGAAAGTAAGCTATATCTTTTCCTTTTTTCTGTTG |
| P77 | TAATAACTCGACGCGGCCGATAACTTCGTATAATGTATGCTATACGAAGTTATACTAGTTCAAGCGTAATCAGGAACGTCGTAG |
| P78 | ATGTACTCTCCTTATGAGGCGCGCCGAGCACGTTTTTTGTTATTTAATTATAATTTCTACTTTTTAGTTATTATTATTATAAATATTTG |
| P79 | CAATACTATATCATTTTATTTTCTTTTCTCTTATATTATATTATTTAATTTTATCTTATTTTTTTTTTTTTTTAAACCTAGGATAACTTCGTATAGCATAC |
